# Wide-ranging and unexpected consequences of altered Pol II catalytic activity in vivo

**DOI:** 10.1101/072785

**Authors:** Indranil Malik, Chenxi Qiu, Thomas Snavely, Craig D. Kaplan

**Affiliations:** Department of Biochemistry and Biophysics, Texas A&M University, College Station, TX 77843, USA

## Abstract

Here we employ a set of RNA Polymerase II (Pol II) activity mutants to determine the consequences of increased or decreased Pol II catalysis on gene expression in *Saccharomyces* cerevisiae. We find that alteration of Pol II catalytic rate, either fast or slow, leads to decreased Pol II occupancy and apparent reduction in elongation rate *in vivo*. However, we also find that determination of elongation rate *in vivo* by chromatin IP can be confounded by the kinetics and conditions of transcriptional shutoff in the assay. We identify promoter and template-specific effects on severity of gene expression defects for both fast and slow Pol II mutants. We show that mRNA half-lives for a reporter gene are increased in both fast and slow Pol II mutant strains and the magnitude of half-life changes correlate both with mutants’ growth and reporter expression defects. Finally, we tested a model that altered Pol II activity sensitizes cells to nucleotide depletion. In contrast to model predictions, mutated Pol II retains normal sensitivity to altered nucleotide levels. Our experiments establish a framework for understanding the diversity of transcription defects derived from altered Pol II activity mutants, essential for their use as probes of transcription mechanisms.

## Introduction

Gene transcription by RNA polymerase II (Pol II) is an essential process and involves three distinct phases: initiation, elongation and termination. Transcription elongation proceeds through an iterative cycle of substrate selection, catalysis of phosphodiester bond formation, and enzyme translocation [Reviewed in (1,2)]. Pausing, backtracking, and arrest of Pol II can occur during elongation. Pol II elongation factors are proposed to promote Pol II elongation by modulating these processes or otherwise enabling Pol II to overcome obstacles. Coordinated with elongating Pol II, several co-transcriptional events occur to control the fate of nascent RNAs and ensure proper gene expression [Reviewed in (3)]. Thus, it is likely that perturbation of Pol II elongation will have multi-faceted effects *in vivo*. To understand the complexity of Pol II activity-mediated control of gene expression it is necessary to understand how alteration of Pol II catalytic activity relates to specific gene expression defects.

Studies from our lab and others have identified several Pol II catalytic mutants that can alter elongation rate *in vitro* (4–6). Based on these mutants’ ability to increase or decrease transcription elongation rate relative to wild type (WT), we term them “gain of function” (GOF) or “loss of function” (LOF) mutants, respectively (4). These classes of mutant confer distinct gene expression profiles, genetic interaction profiles, splicing, and transcription initiation defects *in vivo*, consistent with their having distinct biochemical defects *in vitro* (4,7,8). Most of these mutants reside in a highly conserved, mobile sub-domain of the largest Pol II subunit Rpb1, known as the trigger loop (TL) (4–6,9). The TL is a component of the Pol II catalytic center, and can directly interact with incoming NTPs, undergoing conformational changes to promote rapid catalysis (5,10,11). TL mutants have been shown to affect a number of Pol II biochemical properties including catalysis, substrate selection and transcription fidelity (5,11). In addition, TL mutants have been also shown to affect Pol II translocation, pausing and intrinsic cleavage properties (12–14).

The rate of transcription elongation has likely evolved to facilitate and enhance the efficiency of pre-mRNA processing and maturation [Reviewed in (3)]. Maturation of pre-mRNA requires addition of a 7-methyl guanosine cap at the 5’-end of the transcript, splicing of introns, and addition of a poly(A) tail to the 3’-end of the transcript. Further, the pre-mRNA is uniquely packaged with protein components into a mature mRNA granule, which facilitates export and efficient translation. Impaired processing leads to degradation of pre-mRNAs by nuclear surveillance pathways. Mechanistic coupling of transcription and pre-mRNA processing is achieved through recruitment of factors by C-terminal domain (CTD) and by kinetic competition between transcription and processing [Reviewed in (3)]. Using Pol II catalytic mutants, it has been shown that kinetic competition functions in the efficiency of pre-mRNA splicing (7,15–17). At least one Pol II catalytic mutant has been reported to be defective in 5’-capping, leading to the degradation of transcript by 5’ to 3’ nuclear exonuclease (18). Additionally, a number of findings suggest kinetic competition between transcription termination and elongation in both yeast and human cells (19,20). Recently, a competition-independent pathway has been proposed for termination that occurs through a conformational change to Pol II (21). Furthermore, it has been proposed that overall RNA synthesis rate is connected to mRNA decay rate through feedback between synthesis and degradation (22–26). Growth rate is also proposed to control mRNA decay and overall mRNA abundance, but linkages between RNA synthesis rate, growth, and mRNA decay have not been determined at the molecular level (27,28)(http://biorxiv.org/content/early/2016/03/19/044735).

In order to study Pol II elongation *in vivo*, a number of methods have been implemented that either directly measure apparent Pol II elongation rate or determine indirect consequences of elongation rate (29–32) (for a of list methods and estimated *in vivo* Pol II elongation rates see here (33)). Two methods are generally used to study elongation in yeast, one to study elongation properties and the other to genetically implicate factors in elongation control. The first utilizes chromatin immunoprecipitation (ChIP) to determine Pol II occupancy across a long galactose-inducible gene, *GAL1p∷YLR454w*, either in steady state or after transcription shutoff by addition of glucose (29). Apparent processivity is inferred from comparison of steady state Pol II occupancy for wild type and transcription mutants, while kinetics of the ‘last wave’ of Pol II leaving the template can be used to determine the apparent elongation rate. Indeed, Pol II catalytic mutants and several mutant factors have shown altered apparent *in vivo* elongation rate and apparent processivity defects in this assay (**Supplementary Table 1**). Interpretation of apparent elongation rate differences based on transcriptional shutoff makes assumptions that signaling and kinetics of the shutoff are identical between WT and mutant strains. Similarly, in a second widely used approach, genetic detection of elongation defects through use of nucleotide-depleting drugs makes assumptions that drug effects are identical between WT and mutant strains, and this issue is discussed below.

Nucleotide-depleting drugs, such as mycophenolic acid (MPA), which limits cellular GTP levels by inhibiting IMPDH activity, are assumed to elicit transcription elongation defects by enhancing pausing, arrest, or backtracking due to limitation in substrate levels (1,34–37). Growth sensitivity to MPA for Pol II or presumptive elongation factor mutants has been widely interpreted as a synergistic effect between MPA treatment and impaired elongation due to the mutant. The notion that limiting nucleotide is the major determinant of drug phenotypes was further strengthened by the observation that guanine supplementation suppresses sensitivity to the drug, along with the observation of elongation defects due to drug treatment (29,38). However, it has been shown subsequently that many MPA-sensitive transcription mutants are defective for upregulation of the MPA-resistant IMPDH activity encoded by the *IMD2* gene (39). Under GTP-replete conditions, an upstream transcription start site (TSS) at *IMD2* is used (where transcripts initiate using GTP), generating an unstable transcript that terminates within the *IMD2* promoter (40,41). Upon GTP starvation elicited by MPA treatment, a downstream TSS is utilized (where transcripts initiate using ATP), allowing for expression of functional *IMD2*. Regulation of *IMD2* expression, presumably mediated by this TSS switch, is defective in a wide range of transcription elongation mutants, as well as for mutants that alter Pol II catalytic activity (4,38,42). The mechanism for defects in *IMD2* TSS switching has not been determined for any mutant. An attractive model for Pol II mutant effects at *IMD2* is that Pol II catalytic defects mimic changes to NTP substrate levels, and suggest that the Pol II active site may directly communicate GTP levels to the *IMD2* promoter through initiation efficiency at different transcription start sites.

Here, we present a detailed molecular analysis of how alteration to Pol II catalysis rate through the use of specific *rpb1* alleles, conferring slower or faster catalysis than wild type *in vitro*, affects several facets of transcription and gene expression *in vivo* (a summary of observed mutant effects is in **Supplementary Table 2**). Using the widely used *GAL1*p∷*YLR454w* reporter, we show that both slow and fast Pol II catalytic mutants decrease Pol II occupancy and reporter gene expression *in vivo*. We find that the reporter gene expression defects in slow or fast Pol II mutants negatively correlate with increased reporter mRNA half-lives. Pol II catalytic mutants show genetic interaction with pre-mRNA processing factors, and we present evidence for an mRNA processing defect in fast catalytic mutants. Finally, we have critically evaluated the two major widely used systems for studying transcription elongation in yeast – chromatin IP of the ‘last wave’ of Pol II upon transcription inhibition, and response to GTP starvation. For both assays we uncover underlying biological complexities that differentially affect WT and presumptive elongation mutant cells, confounding interpretation of these assays by simple models. Our results provide a useful framework for future utilization of Pol II catalytic mutants to probe biological processes and gene expression mechanisms.

## MATERIAL AND METHODS

### Yeast strains, plasmid, media and growth

Yeast strains and plasmids used in this study are listed in **Supplementary Table 3 and 4, respectively**.

Yeast media are prepared following standard (43) and previously described protocols (4). Yeast extract (1% w/v; BD), peptone (2% w/v; BD) and 2% bacto-agar (BD), supplemented with adenine (0.15mM) and tryptophan (0.4mM) (Sigma-Aldrich) comprised YP solid medium. YPD plates contained dextrose (2% w/v, VWR), YPRaf plates contained raffinose (2% w/v, Amresco), YP-Raf/Gal plates contained raffinose (2% w/v) plus galactose (1% w/v, Amresco) and YPGal plates contained galactose (2% w/v) as carbon sources. YPRaf, YPGal and YPRaf/Gal plates also contained Antimycin A (1 mg/ml; Sigma-Aldrich). Minimal media plates were prepared with synthetic complete (SC) or ‘Hopkins mix’ with appropriate amino acid(s) dropped out as described in (43), with slight modifications as described in (4). For studies with mycophenolic acid (MPA, Sigma-Aldrich), a stock solution (10 mg/ml, in 100% ethanol) of MPA was added to solid or liquid media to achieve desired concentration. NaOH, HCl and Guanine were added to solid media to achieve desired concentration as indicated. Liquid YPD, YPRaf, YPGal and YPRafGal media are prepared with yeast extract (1% w/v), peptone (2% w/v) and 2% (w/v) carbon source (dextrose, raffinose, galactose or raffinose plus galactose), with no supplementary adenine, tryptophan or Antimycin A. Antimycin A is a standard additive to yeast plates when growth on fermentable carbon sources such as galactose or raffinose is being assayed. Prevention of respiration enforces growth by fermentation. These assays can be performed in the absence of Antimycin A (which is a highly toxic substance), but they become less sensitive as the growth differenced between WT *gal10*∆*56* strains on YPRafGal and mutants that suppress galactose toxicity on YPRafGal are lessened.

Yeast phenotyping assays were performed by serial dilution and spotting onto plates as described earlier (4). Doubling times for the mutants in liquid medium (YPGal) were determined using Tecan plate-reader as described earlier (44), with minor modifications. Overnight grown saturated cultures were diluted to an OD_600_ of ~ 0.1 in fresh YPGal medium and grown in triplicate at 30˚C in a 96-well plate in a Tecan Infinite F200 plate reader under continuous shaking. Data obtained from each plate were considered as a single biological replicate and was analyzed in Graphpad Prism using an exponential-growth fitting function.

### Chromatin immunoprecipitation

All the strains used for ChIP experiments contained a C-terminal epitope tag on the Rpb3 subunit of RNA Pol II (RPB3∷3XFLAG∷KANMX; see strain list). ChIP experiments were performed as described previously (45). Briefly, 100 ml of mid-log phase cells (~1×10^7^ cells/ml) were cross-linked with 1% formaldehyde (final) for 20 min, and then quenched with 15 ml of 2.5 M glycine for 5 min. Cross-linked cells were washed twice with cold 1× TBS buffer at 3500 rpm for 3-4 min at 4˚C using a JS-5.3 rotor (Avanti J-26 XP centrifuge, Beckman-Coulter) and were disrupted by bead beating with glass beads in lysis buffer (0.1 M Tris pH 8.0, glycerol 20%, 1 mM PMSF). Cross-linked cell lysates were subjected to a low speed spin (1500 rpm, 1 min at 4˚C) to remove cell-debris, followed by centrifugation of chromatin pellets, subsequent washing of pellets (twice) with 1 ml FA buffer (50 mM HEPES-KOH pH 7.5, 300 mM NaCl, 1 mM EDTA, 0.1% Triton X-100, 0.01% sodium deoxycholate, 0.1% SDS and 1 mM PMSF) at 14000 rpm for 20 min at 4˚C using F45-30-11 rotor (Centrifuge 5417R, Eppendorf). Chromatin pellets were resuspended in 1 ml of FA buffer and sonicated at 4˚C using a Diagenode Bioruptor (45 cycles – 3 × 15 cycles; 30 sec ON/ 45 sec OFF) to generate ~300-500 bp chromatin fragments (verified on 1% agarose gel). Approximately 100 µl sonicated chromatin was used for each immunoprecipitation (IP) with anti-FLAG antibody (FLAG M2 magnetic beads, Sigma-Aldrich). Surfactant and detergent composition in buffers were changed according to manufacturer’s recommendation for compatibility with M2 FLAG antibody, and all buffers contained 1 mM PMSF. For Pol II occupancy determination, the amount of chromatin used for WT or mutant IPs was normalized by starting cell number and chromatin concentration (estimated by spectrophotometer and agarose gel-lane profile). Magnetically-captured FLAG beads were washed twice with FA buffer, once with FA buffer with 500 mM NaCl, once with wash buffer (10 mM Tris-HCl pH 8.0, 0.25 M LiCl, 0.5% NP-40, 1 mM EDTA) and once with TE. Immunoprecipitated chromatin was eluted by two-step elution at 65˚C, with 100 ul and 150 ul elution buffers (50 mM Tris-HCl pH 8.0, 1 mM EDTA, 1% SDS) for 15 min and 30 min, respectively. Both eluates were pooled and incubated at 65˚C (> 6 hrs) for cross-linking reversal. 10 ul of sonicated chromatin (10% of material added to IP) plus 240 ul of elution buffer were treated identically and served as IP input control. Input or immunoprecipitated DNA was purified by standard phenol-chloroform extraction and ethanol precipitated in presence of pellet paint (MilliporeSigma) or glycoblue (ThermoFisher). Immunoprecipitated DNA and 10% of corresponding input DNA (1:10 diluted in nuclease free water) were used for qPCR with SsoAdvanced or SsoAdvanced Universal SYBR Green supermix (Bio-Rad) using CFX 96 (Bio-Rad). Fold enrichment for target amplicon over non-transcribed region was determined by the ∆∆C_T_ method (46). Primers used for qPCR are listed in **Supplementary Table 5**.

For *in vivo* elongation assays, WT or mutant cells were grown in YPGal to mid-log phase and a pre-glucose shutoff sample was taken for the 0 minute time point. Then glucose (4% final) was added to inhibit transcription and aliquots were removed after 2, 4, 6 and 8 minutes (optional longer time points were taken for some strains). Alternatively, after isolation of 0 min sample as above, remainder of culture was centrifuged and washed with SC medium lacking carbon source as described earlier (19), then inoculated in YPD (4% glucose) media or SC media lacking carbon source to isolate glucose shutoff samples or galactose-depletion samples, respectively, at indicated time points. Formaldehyde cross-linking, chromatin preparation and subsequent steps were performed as described above.

### RNA isolation, Northern blotting

For gene expression analysis, RNAs were isolated from mid-log phase cells grown in appropriate medium as described in the main text. Post shutoff samples were collected by a quick centrifugation for 1 min and immediate freezing of the cell pellets by placing at -80C. Centrifugation time was included while calculating shutoff time. Total RNA was purified using hot phenol-chloroform method as described previously (47).

Northern blotting was performed essentially as described in the manual for GeneScreen hybridization membranes (Perkin-Elmer) with minor modifications as described earlier (4). In brief, 20 µg of total RNA, treated with Glyoxal sample-load dye (Ambion), was separated on 1% agarose gel and transferred to membrane by capillary blotting. Pre-hybridization solution contained 50% formamide, 5× Denhardt’s solution, 10% Dextran sulfate, 1 M NaCl, 50 mM Tris-HCl pH 7.5, 0.1% sodium pyrophosphate, 0.1% SDS and 500 µg/ml denatured salmon sperm DNA. PCR generated DNA double-stranded probes for northern blots were radiolabeled with ^32^P-dATP using the Decaprime II kit (Ambion) according to manufacturer’s instructions. After overnight hybridization of the blot at 42˚C, washes were done twice each in 2× SSC for 15 minutes at 42˚C, in 5× SSC with 0.5% SDS for 30 minutes at 65˚C, and in 0.2× SSC for 30 minutes at room temperature. Blots were visualized by phosphorimaging (Bio-Rad or GE Healthcare) and quantified using Quantity One (Bio-Rad).

Northern blotting for mapping the termination window of the pre-processed *snR33* transcript was performed essentially as described in (19,48) with minor modifications. Briefly, 5-8 µg of RNA were separated on 6% polyacrylamide-7M urea gel. RNAs were transferred from polyacrylamide gel to a membrane (GeneScreen Plus, PerkinElmer) with a Bio-Rad Trans-Blot apparatus at a setting of 45W for 1.5 hrs. RNAs were cross-linked to the membrane by UV light. Pre-hybridization of the membrane, probe synthesis and membranes-hybridization were performed as described in (48). Membranes were washed twice with low stringency wash buffer (0.1× SSC, 0.1% SDS) and visualized by phosphorimaging (Bio-Rad or GE Healthcare). Lane traces were determined for each sample using ImageQuant (GE Healthcare).

### Primer extension

For primer extension (PE) analysis, RNA was isolated from mid-log phase cells grown in appropriate media, optionally treated with desired concentration of MPA for indicated time periods. Primer extension analysis was done essentially as described earlier (49) with modification described in (4). Briefly, 30 µg of total RNA was annealed with ^32^P end-labeled oligo. M-MLV Reverse Transcriptase (Fermentas) was used for reverse transcription, in the presence of RNase Inhibitor (Fermentas). Primer extension products were ethanol precipitated overnight and separated on 8% polyacrylamide gels (19:1 acrylamide:bisacrylamide, Bio-Rad) containing 1× TBE and 7M urea. PE gels were visualized by phosphorimaging (Bio-Rad or GE Healthcare) and quantified using Image Lab software (Bio-Rad). For *IMD2* TSS annotation we have considered ‘A’ of the start codon (ATG) as +1, so that bases upstream start at −1, as in our previous publication (4).

### Microscopy and image analysis

Mig1p-GFP tagged strain was made by integrating GFP C-terminal tag at the genomic locus (see strain description). Microscopy was performed as described previously (50), with modifications. Briefly, cells grown overnight in SC medium (2% galactose) at 30˚C, diluted in fresh SC media and grown till mid-log at 30˚C before microscopy. Perfusion chamber gasket (ThermoFisher, 4 chamber: 19 mm × 6 mm) was used for changing medium. Chamber was treated with Con A (2 mg/ml, MP Biomedicals) for 10-15 min, and then cells were injected into the chamber and allowed 10-15 min to adhere. Medium exchange from SC (2% galactose) to SC (2% galactose) + glucose (4% final) was done by pipetting quickly, while the chamber is fixed on the microscope stage. Pre-glucose sample was considered as time 0 and glucose repression time points were taken immediately after exchange of the medium.

For Mig1p-GFP nuclear localization kinetics, microscopy was performed with an inverted epifluorescence microscope (Ti-E, Nikon, Tokyo, Japan) using a 100× objective (Plan Fluo, NA 1.40, oil immersion) and standard filter sets. Images were acquired using a cooled EMCCD camera (iXon3 897, Andor, Belfast, United Kingdom). For delayed Mig1p response (longer time points), microscopy was performed with a Nikon Ti-E microscope equipped with a CFI Plan Apo lambda DM 100x objective and a Prior Scientific Lumen 200 Illumination system. All images were acquired using NIS Element software and data analysis was done using Quantity One (Bio-Rad). Ratio of nuclear/cytoplasmic GFP intensity was calculated as [(nuclear GFP intensity – background) / (cytoplasmic GFP intensity – background)]. Obtained values for glucose repression time points were normalized to pre-glucose (time 0, t0) value. Non-responding cells were quantified similarly, except for the fact that there was no visible nuclear foci, thus a random, central area was selected and measured for fluorescence intensity. Position and area of the measurement was kept identical for all the time points for the same cell. We reasoned that a non-responding cell would not have a distinct change over time in nuclear/cytoplasmic GFP intensity, hence quantifying a random area as nucleus should not have an overall affect on the interpretation of the data.

## RESULT

### Pol II catalytic mutants decrease steady state Pol II occupancy and apparent processivity

To explore *in vivo* consequences of altered Pol II activity, we examined a subset of Pol II active site mutants in the largest subunit Rpb1. These mutants were previously characterized biochemically, molecularly, and genetically as falling into two classes (4,5,8,9). These two classes of Pol II mutants appear to derive from either increase (“GOF”) or decrease (“LOF”) in Pol II catalytic activity, as determined using *in vitro* transcription. First, we examined Pol II occupancy by chromatin IP over an inducible long reporter gene, *GAL1p∷YLR454w* (Figure 1A and (29)). We observed decreased overall polymerase occupancy for both GOF and LOF mutants compared to WT (Figure 1B). Among the mutants tested, H1085Y (strong LOF) and G1097D (strong GOF) showed the most severe defects in Pol II occupancy, consistent with their severe growth defects *in vivo* (4). When we normalized the Pol II fold-enrichment over the 5’ end of the gene body (“1kb” amplicon) followed by normalization of mutant values to WT, catalytic mutants show selective decrease in occupancy over the 3’ end of the reporter (most obvious for G1097D) (Figure 1C). Decreased steady state Pol II occupancy at the 3’ end is consistent with a processivity defect, wherein Pol II that begins synthesis is unable to complete the transcript, leading to polar effects on occupancy from 5’ to 3’. Generally, greater Pol II occupancy over 5’ ends of transcription units relative to 3’ ends are interpreted as processivity defects when observed (29); however, a faster elongation rate over a gene’s 3’ end would lead to a similar phenotype. Because differential elongation rates over different parts of genes has been observed in mammalian cells (51–53), we are careful to assert that what we observe is differential apparent processivity for Pol II mutants as ChIP is unable to distinguish between the two possibilities.

**Figure 1.**
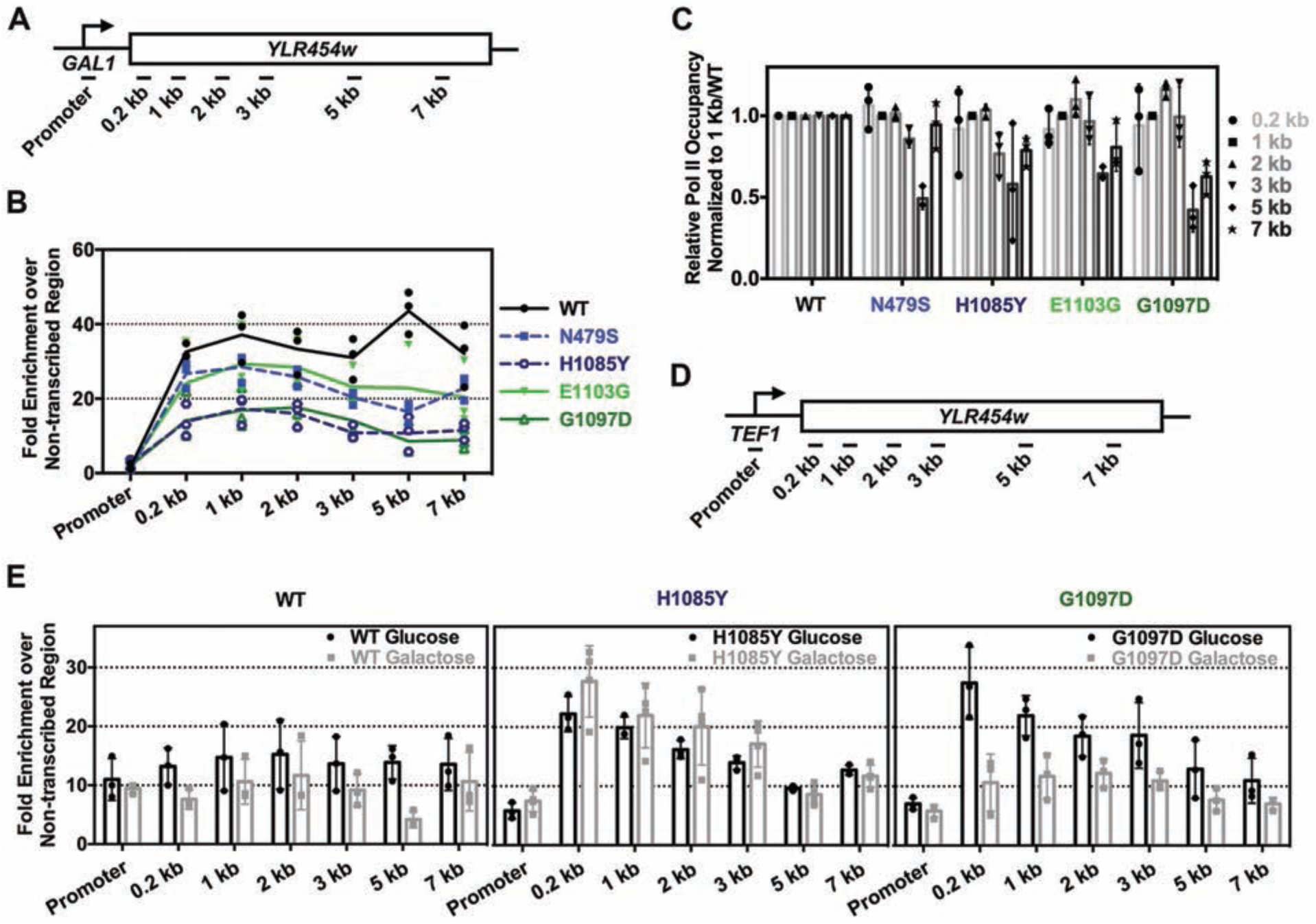
**Pol II catalytic mutants decrease Pol II occupancy and show apparent processivity defects.** (A) Schematic of the galactose inducible reporter *GAL1*p∷*YLR454w* annotated with positions of PCR amplicons used for ChIP experiments. Promoter amplicon derived from primers specific to the *kanmx∷GAL1p* integrated promoter. (B) Pol II occupancy over *GAL1p∷YLR454w* reporter is decreased for both GOF and LOF mutants compared to WT. Line graph shows steady-state Pol II occupancy for WT (black line), GOF (green lines) and LOF (blue dashed lines) mutants under galactose induction. Green and blue color-coding is used to annotate GOF and LOF mutants, respectively. This color-coding is used throughout. (C) Pol II catalytic mutants show apparent 3’ processivity defects. Apparent Pol II processivity defects determined for each mutant by normalizing mutant ChIP signal to signal at 1kb followed by normalization to WT. (D) Schematic of constitutively expressed reporter gene *TEF1p∷YLR454w* annotated with positions of PCR amplicons used for ChIP experiments. Promoter amplicon derived from primers specific to the *kanmx∷TEF1p* integrated promoter. (E) Carbon source may differentially affect Pol II mutants’ occupancy. Comparison of steady-state Pol II occupancy at *TEF1p∷YLR454w* for WT and Pol II catalytic mutants grown in galactose - or glucose-containing medium. Individual data points from at least three biological repeats are shown with bars showing average +/− the standard deviation of the mean.

Additional factors may determine Pol II occupancy levels over genes. For example, Pol II occupancy over a template may be gene class-dependent or regulon-specific, given that external perturbations such as changes in temperature or carbon source have been shown to affect transcription (54,55). Therefore, we examined Pol II occupancy over the same reporter driven by a constitutive promoter (*TEF1p∷YLR454w*) with strains grown in glucose or galactose for the most severely defective Pol II LOF and GOF mutants (Figure 1D and E). We observed an increased Pol II occupancy for H1085Y in galactose relative to glucose, with the converse for G1097D, compared to WT (Figure 1E), although a 3′ occupancy defect was still apparent for both mutants. WT Pol II showed more subtle changes in occupancy in glucose relative to galactose. Furthermore, the G1097D 3′ end was enhanced in glucose relative to galactose, suggesting growth conditional effects on transcription elongation.

### Pol II catalytic mutants show decreases in reporter gene expression that are exacerbated by promoter strength and template length

Defective initiation or elongation might lead to decreased Pol II occupancy (56). Pol II activity mutants do perturb initiation as most or all show defects in TSS selection (7,8). Mutants affecting Pol II elongation have also been shown to affect reporter gene expression depending on length of the transcription unit driven by identical promoters, using the so-called “GLAM” assay (32,57). Therefore, we determined if Pol II catalytic mutants show gene expression defects based on gene length and/or promoter strength. We measured Pol II catalytic mutants’ effects on expression of *YLR454w* reporters in two different promoter contexts, driven by the strong, inducible GAL1 promoter and the constitutive TEF1 promoter by Northern blotting (Figure 2A). Additionally, we also measured expressions of the endogenous *GAL1* and *TEF1* genes for comparison with *YLR454w* driven by the same promoters. Pol II mutants confer the strongest effects on *GAL1*p∷*YLR454w* expression compared to WT, with H1085Y and G1097D having the most severe effects (Figure 2A). Furthermore, we observed greater gene expression defects when examining the highly expressed *GAL1*p∷*YLR454w* relative to the more lowly expressed *TEF1*p∷*YLR454w*. We also found that expression defects for the *YLR454w* reporters were greater than defects observed for the native genes (shorter in length) under control of their respective promoters (Figure 2A and 2B). These results are consistent with previous observations showing Pol II occupancy - and gene-length-dependent effects on reporter expression for several elongation mutants (32,56,57). Given reduced Pol II occupancy and gene expression, it is likely that initiation defects might also be in play. When we examined the kinetics of activation for *GAL1p∷YLR454w* or *GAL1*, we found that activation of expression was greatly delayed in Pol II mutant strains, consistent with defects in activation or initiation (Figure 2C).

**Figure 2.**
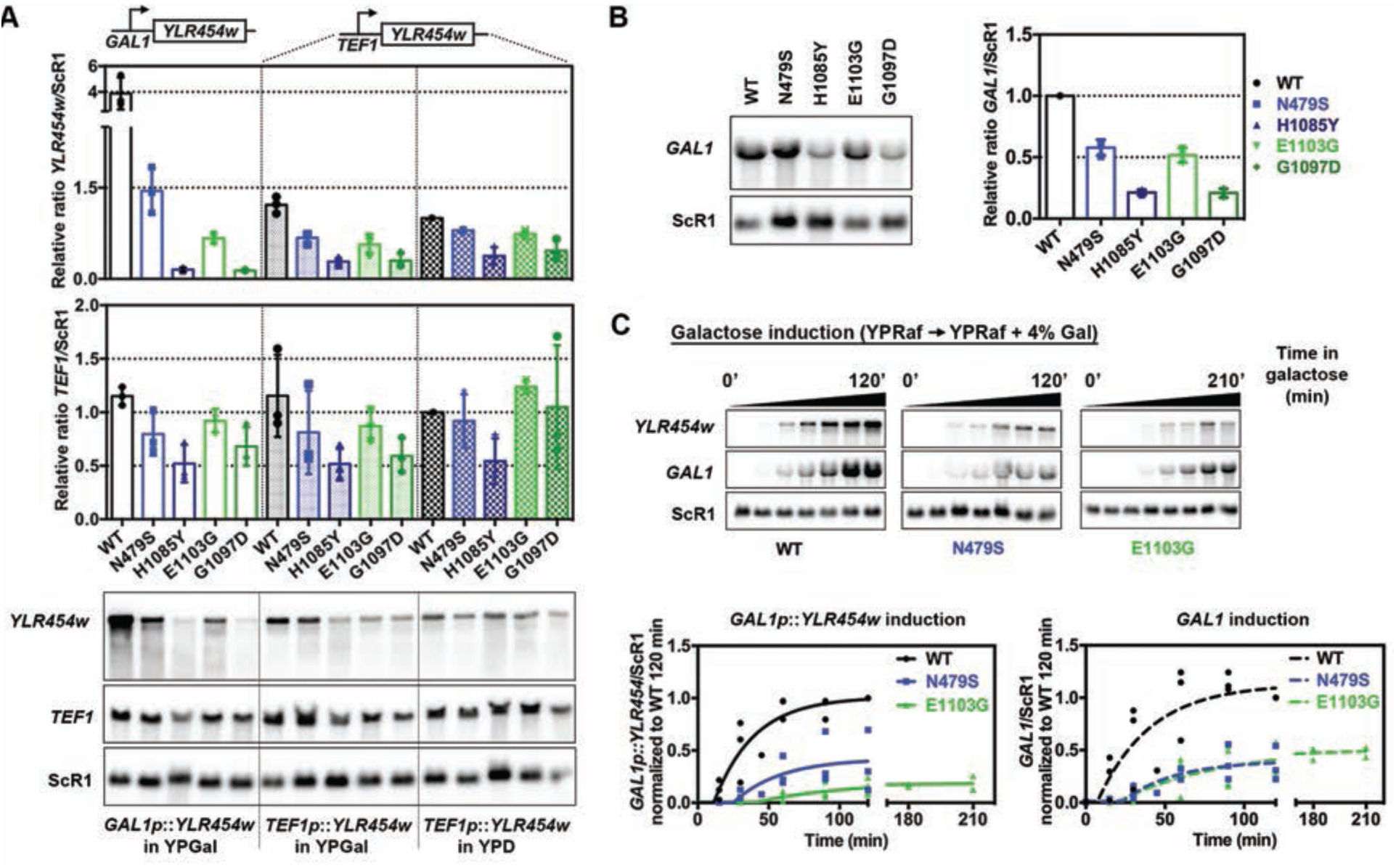
**Pol II catalytic mutants generally decrease in vivo gene expression.** (A) Pol II mutant effects on gene expression are exacerbated with increasing promoter strength. Steady-state RNA levels of reporter genes used for Pol II occupancy experiments (*GAL1p∷YLR454w* and *TEF1p∷YLR454w*) and endogenous *TEF1* levels from cells grown in galactose - or glucose-containing medium as indicated. Values were normalized to WT *TEF1p∷YLR454w* expression level in YPD. The most severe expression defects are evident at *GAL1p∷YLR454w* for both GOF and LOF mutants compared to WT. Individual data points from at least three biological repeats are shown with error bars indicating average +/−standard deviation of the mean. (B) Endogenous *GAL1* mRNA expression level is decreased in Pol II catalytic mutant strains compared to WT, yet *GAL1* expression defects in mutants are less severe than *GAL1p∷YLR454w* expression defects as showed in A. Values were normalized to WT *GAL1* mRNA level. Error bars as in A. (C) Pol II catalytic mutants delay induction of the *GAL1* promoter. Time courses showing induction of *GAL1p∷YLR454w* and endogenous *GAL1* mRNA in WT and Pol II catalytic mutants. Overnight grown cells were inoculated into fresh YPRaf medium and grown until mid-log phase at 30˚C, subsequently 4% galactose (final) was added to induce *GAL* gene expression. RNAs isolated prior (time 0) and after galactose addition were used for Northern blotting to determine accumulation of mature *GAL1p∷YLR454w* and endogenous *GAL1*. Data normalized to WT 120 min value and plotted using non-linear regression using GraphPad Prism. Individual data points from at least three biological repeats are shown.

### Pol II catalytic mutants show allele-specific genetic interactions with pre-mRNA processing factors

We observed greater expression defects for the Pol II mutants compared to occupancy defects over the same reporter *GAL1p*∷*YLR454w* (compare Figure 1B with Figure 2A). These results suggested that there could be additional defects beside Pol II initiation or elongation defects contributing to decreased abundance of the reporter mRNA in the mutants. As the abundance of cellular mRNAs is determined by synthesis and degradation rates, we reasoned that overall expression defects observed for Pol II mutants might stem from changes to mRNA stability and/or RNA processing. To start, we determined if Pol II catalytic mutants show *GAL1*p∷*YLR454w* processing defects. We took a two-pronged approach to address processing defects. First, we examined genetic interactions between Pol II mutants and pre-mRNA processing/degradation factors; second, we asked if blocking pre-mRNA degradation pathways could rescue gene expression defects of Pol II mutants.

We reasoned that an mRNA capping defect or premature termination could lead to degradation of pre-mRNA by nuclear 5′ or 3′-exonucleases, respectively, resulting in decreased abundance of *GAL1*p∷*YLR454w*. Thus, we tested if Pol II mutants showed genetic interactions with known nuclear exonucleases that function in surveillance. Deletion of Rrp6, a component of the 3′-exonuclease exosome complex, showed a slight negative genetic interaction with Pol II GOF mutants (Figure 3A). Inactivation of Rat1 and Xrn1, two 5′-exonucleases, showed strong genetic interactions with GOF mutants (Figure 3A). Rat1 is essential and functions in termination of Pol II at protein-coding genes (58,59). Pol II LOF mutants partially suppressed the Ts^-^ phenotype of *rat1-1*, a conditionally viable allele of *RAT1*. In contrast, Pol II GOF alleles showed synthetic sick interactions with *rat1-1*, consistent with prior findings and model proposed for kinetic competition between termination and elongation (19,20,60). Rat1 works with an activating partner Rai1, which processes partially capped mRNA allowing Rat1 access (61). Growth defects of *rai1∆* were slightly suppressed by both LOF and GOF mutants at 37˚C, while *dxo1∆*, a recently described nuclear 5′-exonuclase (62), showed no strong genetic interactions with Pol II alleles (Figure 3A).

**Figure 3.**
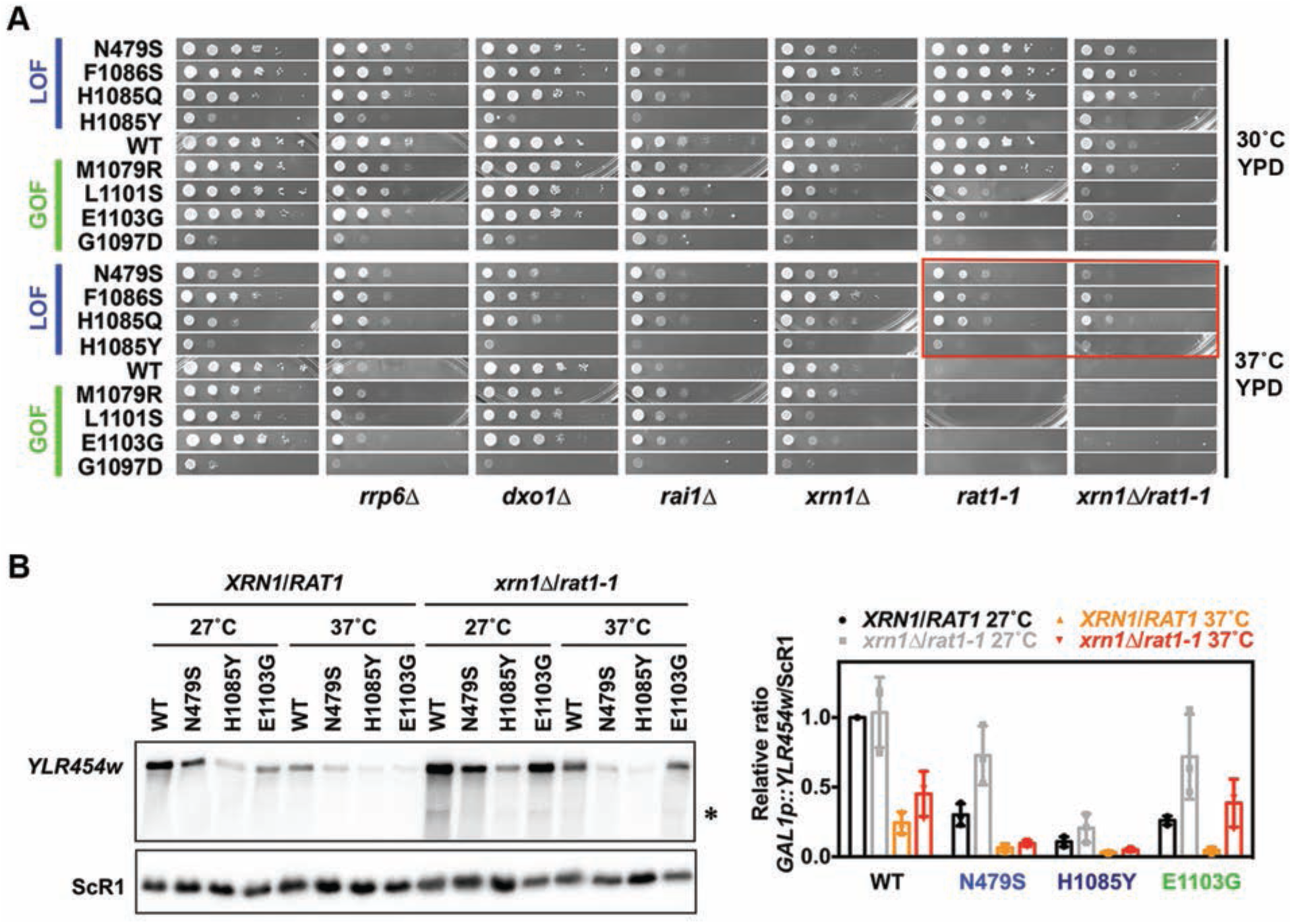
**Allele-specific genetic interactions between Pol II catalytic mutants and pre-mRNA processing factors.** (A) Pol II catalytic mutants show allele-specific genetic interactions with 5’ exonuclease mutants. 10-fold serial dilutions of saturated cultures of Pol II catalytic mutants alone or in combination with pre-mRNA processing factor mutants plated on YPD for comparison of growth at 30˚C and 37˚C. Suppression of *rat1-1* lethality at restrictive temperature is highlighted in red box. (B) Inactivation of both Xrn1 and Rat1 rescue *GAL1*p∷*YLR454w* expression in the Pol II GOF mutant E1103G. (Left) Representative gel showing *GAL1p∷YLR454w* reporter expression in Pol II catalytic mutants in WT and *xrn1*∆/*rat1-1* mutant background. Overnight grown cells were inoculated in fresh YPRaf/Gal media to amplify until mid-log at permissive temperature (27˚C) then shifted to restrictive temperature (37˚C) to inactivate Rat1-1p. RNAs isolated from the half of the culture before shifting temperature used as 27˚C sample. The other half was washed and resuspended in pre-warmed YPRaf/Gal media to grow for another 2 hrs at 37˚C to isolate 37˚C sample for Northern blotting. Asterisk indicates an internal cryptic *GAL1*p∷*YLR454w* transcript in *xrn1*∆/*rat1-1* mutant background, as reported earlier for an *spt16* mutant (90). (Right) Quantification of *xrn1*∆/*rat1-1* effects. Values were normalized to WT *GAL1*p∷*YLR454w* mRNA level (*XRN1*/*RAT1*) at 27˚C. Data shown are average of three biological repeats with error bars representing standard deviation of the mean.

We then asked if the mRNA processing mutants that showed genetic interactions with Pol II mutants were able to modulate Pol II alleles’ gene expression defects. As predicted from the lack of strong genetic interactions, deletion of Rrp6 did not rescue full-length *GAL1*p∷*YLR454w* expression level significantly; furthermore the nuclear exosome would be expected to mainly degrade prematurely-terminated transcripts and not-full length ones (**Supplementary Figure 1A**). Deletion of *XRN1* slightly rescued *GAL1*p∷*YLR454w* expression level in Pol II GOF mutant E1103G (**Supplementary Figure 1A**). Inactivation of Rat1 (at the restrictive temperature, 37˚C) severely decreased *GAL1*p∷*YLR454w* expression levels in both WT and catalytic mutants possibly due the *rat1-1* growth defect, although relative levels of *GAL1*p∷*YLR454w* in WT and Pol II mutants didn’t differ greatly between permissive (27˚C) and restrictive temperature (37˚C) (**Supplementary Figure 1B**). We found that *xrn1*∆/*rat1-1* double mutant strains were extremely sick on YPGal even at permissive temperature (**Supplementary Figure 1C**). Hence, for determination of *GAL1*p∷*YLR454w* expression level we used YPRaf/Gal liquid medium, which allows growth of Pol II mutants in the *xrn1*∆/*rat1-1* background. Intriguingly, inactivation of both Xrn1 and Rat1 together completely rescued the *GAL1*p∷*YLR454w* expression level in E1103G (*xrn1*∆/*rat1-1*) compared to WT (*xrn1*∆/*rat1-1*) at 37˚C (Figure 3B), suggesting possible 5′ end processing-defective or 5′-cap defective transcripts in the GOF mutant E1103G are stabilized by inactivation of Xrn1 and Rat1. Taken together these genetic interaction results suggest Pol II GOF mutants may have 5’ end processing or capping defects, rendering transcripts sensitive to Rat1/Xrn1.

### Both slow and fast Pol II catalytic mutants confer increased mRNA stability to a reporter gene

We reasoned that in addition to myriad Pol II transcription defects and pre-mRNA processing defects discussed above, compromised mRNA stability of the *GAL1*p∷*YLR454w* transcript may also contribute to its lower abundance in Pol II catalytic mutants. In order to determine if Pol II mutants alter mRNA stability, we performed transcriptional shutoff followed by measurement of *GAL1*p∷*YLR454w* mRNA over a time course by Northern blotting. mRNA half-lives were determined by fitting decay curves with a lag followed by exponential decay (**Supplementary Figure 2A**). We found that all GOF and LOF mutants conferred an increase in both the *YLR454w* half-life and lag periods prior to exponential decay relative to WT (Figure 4A, **Supplementary Figure 2B-H**). Further, the increases in mRNA half-lives positively correlated with the overall decrease in expression level of the reporter in the mutants (Figure 4B, **Supplementary Figure 2I**). These results suggest a number of conclusions. First, they indicate defects in mRNA synthesis rates for all tested Pol II mutants, as steady state expression levels were reduced relative to wild type even though we found that the *GAL1-YLR454w* mRNA showed increased stability. Second, longer lag periods prior to exponential decay may indicate Pol II mutants confer a delayed response for transcriptional shutoff, slower elongation rate on this template, or delayed mRNA export (addressed below). Such lags have been observed in genome-wide experiments for mRNA stability where transcription inhibitor blocks initiation but elongation is allowed to proceed (63). The lag can result from the time it takes polymerases that have initiated prior to the block to finish synthesis of their messages. Given that mRNA decay is strongly coupled to translation, and translational demand is coupled to growth rate, it may be difficult to deconvolute Pol II mutant effects on mRNA decay through changes to growth rates versus direct effects on mRNA decay. Indeed, we observed that mRNA half-lives positively correlated with strain doubling times (Figure 4C, **Supplementary Figure 2J**).

**Figure 4.**
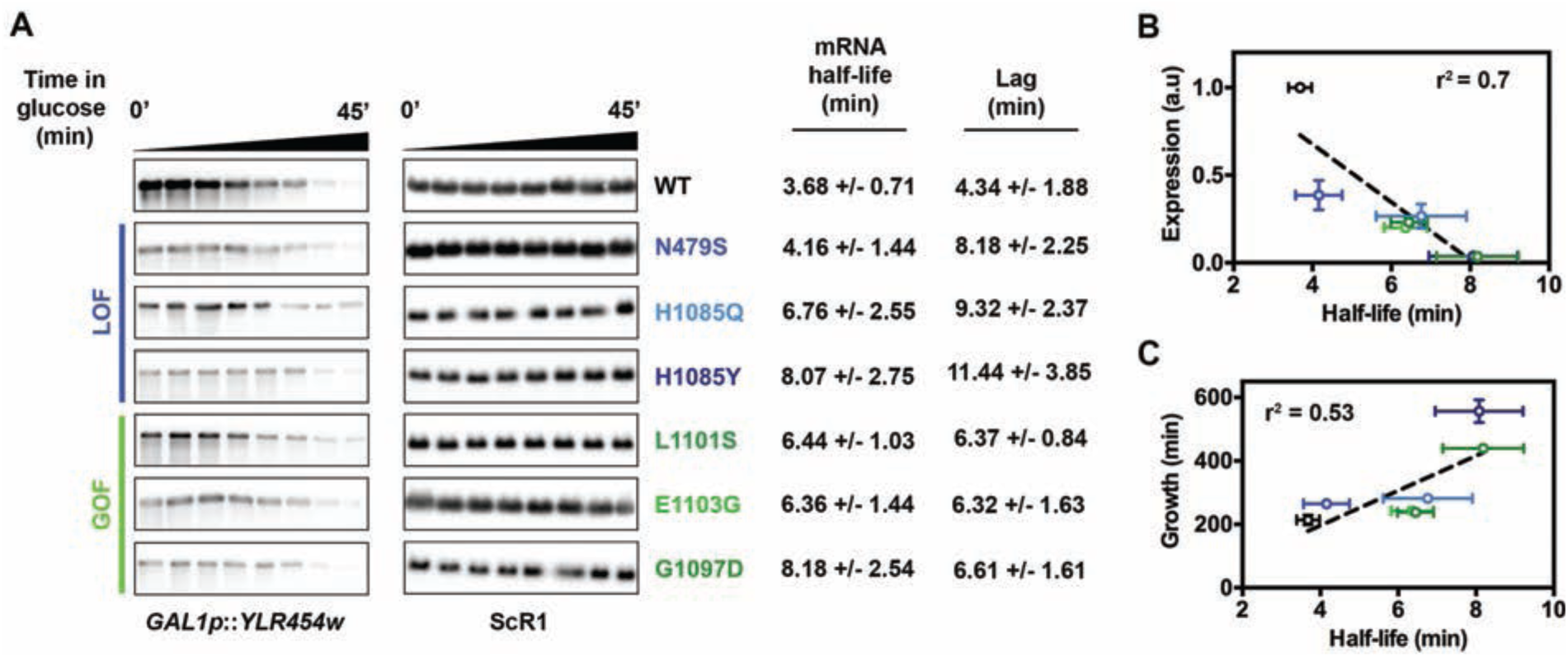
**Pol II catalytic mutants increase mRNA half-life of the reporter.** (A) Both GOF and LOF mutants show reduced degradation rate of *YLR454w* transcript compared to WT. Representative blots showing *GAL1p∷YLR454w* mRNA decay, determined by glucose shutoff. Overnight grown cells were inoculated in fresh YPGal media to grow till mid-log at 30˚C. RNAs isolated from pre-glucose addition (time 0) and post-glucose addition time points were used for Northern blotting. mRNA half-lives and lags were determined using plateau followed by one-phase decay curve fitting in GraphPad Prism (see **Supplementary Figure 2A**). Values indicate average of a minimum of five biological repeats +/−standard deviation of the mean. (B and C) Correlation of *GAL1p∷YLR454w* half-life in different mutants with the expression level (Figure 2A and **Supplementary Figure 2I**) and with the mutants’ growth defects (**Supplementary Figure 2H**) determined by linear regression using GraphPad Prism.

### Slow and fast Pol II catalytic mutants exhibit slower apparent *in vivo* elongation in a commonly used ChIP assay

Our observation of longer mRNA decay lag times than WT for all Pol II mutants tested, both slow and fast, was unexpected under the presumption that Pol II mutants that are fast elongaters *in vitro* are also fast *in vivo*. In our mRNA decay experiments, longer lag times prior to exponential decay for LOF mutants were consistent with delayed clearance of the template and predicted slow elongation kinetics. Conversely, GOF mutants would be predicted to run off the template more quickly. In contrast to this prediction, GOF mutants also showed longer lag times prior to exponential decay (Figure 4A). One GOF mutant tested here, E1103G, was previously suggested to be faster than WT *in vivo* based on chromatin IP analysis of the same *GAL1*p∷*YLR454w* reporter used here (19). Therefore, we examined elongation of E1103G and additional Pol II alleles *in vivo* using this commonly used ChIP assay. In this assay, transcription of *GAL1*p∷*YLR454w* is shutoff by addition of glucose to the medium, and the kinetics of the last wave of Pol II transcription clearing the gene is measured using ChIP (Figure 5A and 5B) (29). As predicted, LOF mutants N479S and H1085Y showed extensive delay in Pol II run-off kinetics following the transcriptional shut-off (Figure 5C and D). The apparent *in vivo* elongation rate of H1085Y was slower than N479S, which is consistent with its stronger growth defects and *in vivo* phenotypes relative to N479S. Surprisingly, both test GOF mutants, E1103G and G1097D, also showed reduced kinetics of Pol II runoff subsequent to glucose addition (Figure 5E). This result was unexpected, especially as E1103G was reported to have a faster apparent elongation rate than WT in a version of this assay (19). Furthermore, analogous substitution in human POLR2A E1126G confers a slightly faster than WT elongation rate *in vivo* (15).

**Figure 5.**
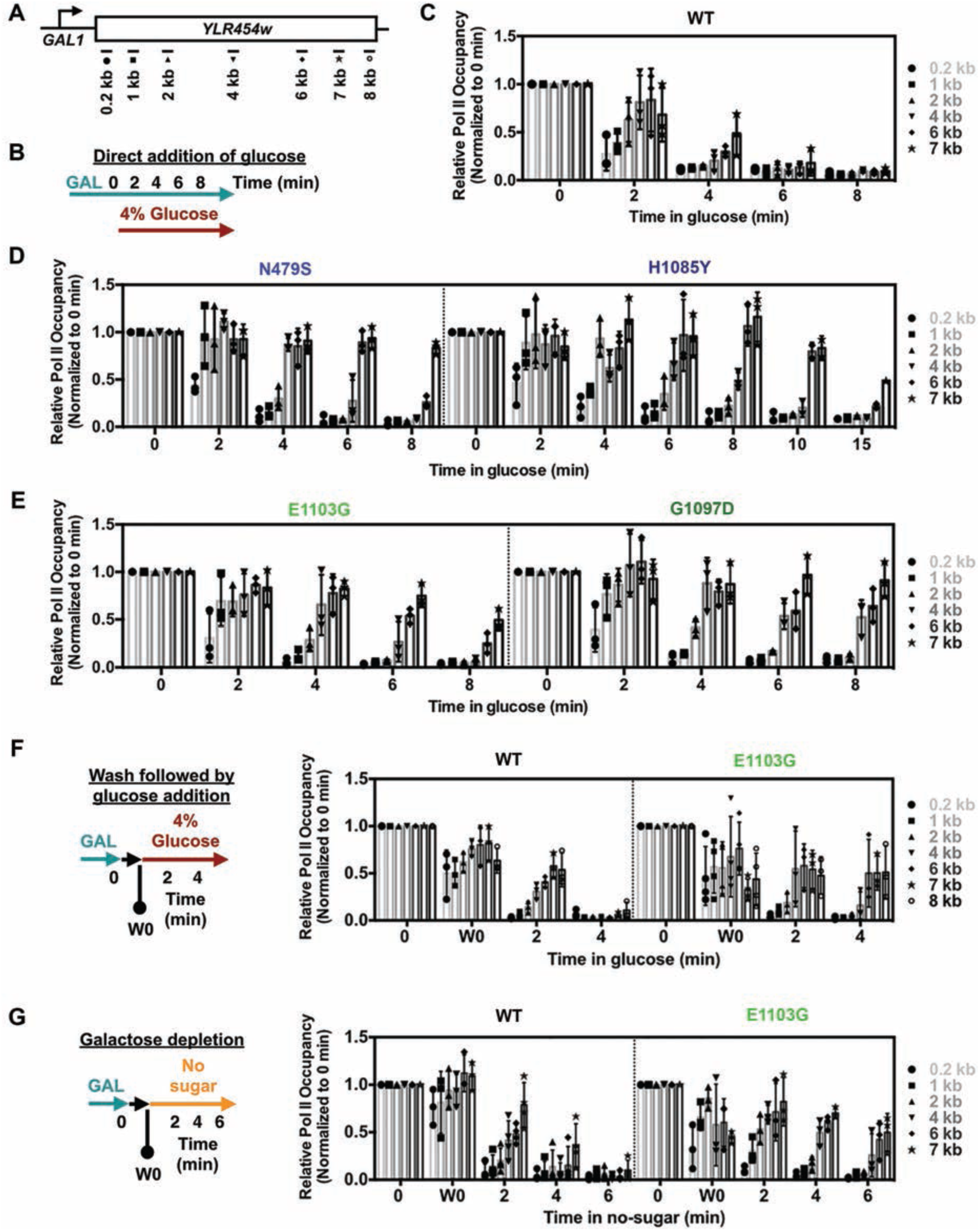
**Pol II catalytic mutants show slower in vivo elongation rate in glucose shutoff assay.** (A) Schematic of the *GAL1p∷YLR454w* with positions of PCR amplicons used for ChIP experiments. (B) Schematic of the regular glucose shutoff experiment used in C-D. *In vivo* apparent elongation rate for WT (C), LOF (D; N479S and H1085Y) and GOF (E; E1103G and G1097D) catalytic mutants determined by ChIP upon glucose shutoff of transcription by direct addition of 4% glucose (final) to the mid-log grown culture in YPGal at 30˚C. Values were normalized to pre-glucose addition (0 min) and error bars represent standard deviation of the mean for at least three independent experiments. For H1085Y, longer time point (10 and 15 min) values obtained from two repeats, with error bars indicating the range of the two experiments. (F) Glucose shutoff assay to compare apparent *in vivo* elongation rate between WT and fast catalytic mutant (E1103G) following Hazelbaker et al., protocol (19). Pre-glucose addition sample (0 min) was isolated as described in Figure 5A; subsequently cells were washed in synthetic complete media lacking carbon source and inoculated in YPD (4% dextrose) to shutoff the transcription. One wash 0 (W0 min) sample was isolated after the washing and before shutoff to determine the effect of washing. Values were normalized to pre-glucose addition (0 min) and error bars represent standard deviation of the mean of at least three independent repeats. (G) Galactose depletion to determine apparent elongation rate in WT and fast catalytic mutant (E1103G). Pre-glucose (0 min) and wash 0 (W0 min) samples were taken as described in Fig 5G, followed by inoculation of cells into synthetic media lacking any carbon to incur transcriptional shutoff in absence of any sugar. Values were normalized to pre-glucose addition (0 min) and error bars represent standard deviation of the mean for at least three independent repeats.

Given the discrepancy between our results and those of Hazelbaker et al (19), we felt it was essential to closely examine Pol II mutant behavior in the elongation assay and identify if there is any hidden or confounding variable. We first extensively validated our GOF mutant strains in three ways – i. confirmation of introduced mutations by sequencing, ii. confirmation of transcriptional growth phenotypes as described in (4), and iii. confirmation of mutants’ ability to shift transcriptional start sites (TSSs) at the *ADH1* promoter as described in (4,7,8) (**Supplementary Figure 3A**). In all cases, results were consistent with strains showing expected phenotypes, leading us to speculate that there might be confounding variable(s) in the ChIP elongation assay itself. While we employed the most commonly used method to shutoff *GAL* transcription by direct addition of glucose to the media (29), in Hazelbaker et al (19), cells were washed in the absence of any carbon source prior to addition of glucose. We repeated the method of (19) with the addition of a critical control, taking a time point just post wash (W0) to determine how the wash step affected Pol II occupancy. In our hands, GOF E1103G was still slower exiting the template than WT (Figure 5F), consistent with both our direct glucose shutoff without washing, and our measurement of increased lag in mRNA decay experiments (Figure 5E and 4A). Additionally, we observed two more phenomena during the wash. First, the W0 time point showed evidence of transcriptional shutoff for both WT and E1103G, presumably due to galactose depletion (see below for further discussion). We confirmed that Pol II run off was not due to delayed formaldehyde cross-linking, as we did not detect any difference in Pol II occupancy when crosslinking was simultaneous with the addition of glucose (**Supplementary Figure 3B**). Second, GOF E1103G showed a faster loss of Pol II over the 3’ end of *GAL1p∷YLR454w* early in the time course (W0) that was not observed in the direct glucose shutoff assay (compare Figure 5E with 5F). Together, these results indicate that the mode of transcription inhibition could alter Pol II transcriptional properties in unexpected ways, additionally confounding ability to estimate apparent Pol II elongation rate.

The absence of galactose itself is a distinct mechanism of transcription shutoff for *GAL* genes. Depletion of galactose will lead to reestablishment of Gal80-mediated repression of Gal4, the critical activator of *GAL* transcription. To examine this mode of transcription shutoff further, without the mixing of two distinct modes of repression, we performed a galactose starvation/depletion time course where no sugar was added. We found similar 3’ specific runoff in E1103G early in the time course (W0) when a wash was employed (Figure 5G). As Pol II runs into the gene 3’ end at later time points (presumably), the effect diminishes. Additionally, in this iteration of the experiment there appears to be faster loss of E1103G than WT from the 5’ end of the reporter early, though E1103G is maintained much longer over the 3’ end later in the time course (Figure 5G). Taken together, we find that Pol II GOF mutants, where tested, appear to be retained longer on the template, when examined in the *GAL1*p*∷YLR454w* system. We observe this under three distinct transcription shutoff conditions and in all cases our results contradict previously published results for Pol II GOF E1103G. However, our studies suggest that E1103G effects on the 3’ end of *GAL1*p*∷YLR454w* upon galactose starvation (during wash) and unintended variability in wash time between strains could account for the observed differences in our experiments and previous observations.

Finally, we examined alternative *in vivo* evidence for putative fast and slow elongation of Pol II GOF and LOF mutants. We analyzed how Pol II mutants altered termination at *SNR33*. It has been proposed previously that kinetic competition between elongating Pol II and Sen1 helicase determines the termination of the snR33 transcript (19). Examination of snR33 intermediate transcripts prior to complete processing allows detection of the snR33 termination window (Figure 6A). Both GOF mutants E1103G and G1097D increase the average length of pre-processed snR33, which is in agreement with previous observations for E1103G (Figure 6B and (19)). Conversely, LOF mutants N479S and H1085Y decreased the average length of pre-processed snR33 (Figure 6B). We also observed alternative mature snR33 transcripts longer or shorter in length for GOF and LOF mutants, respectively, consistent with increased upstream or downstream utilization of transcription start site (TSS) in these mutants as predicted for alterations in catalytic rate.

**Figure 6.**
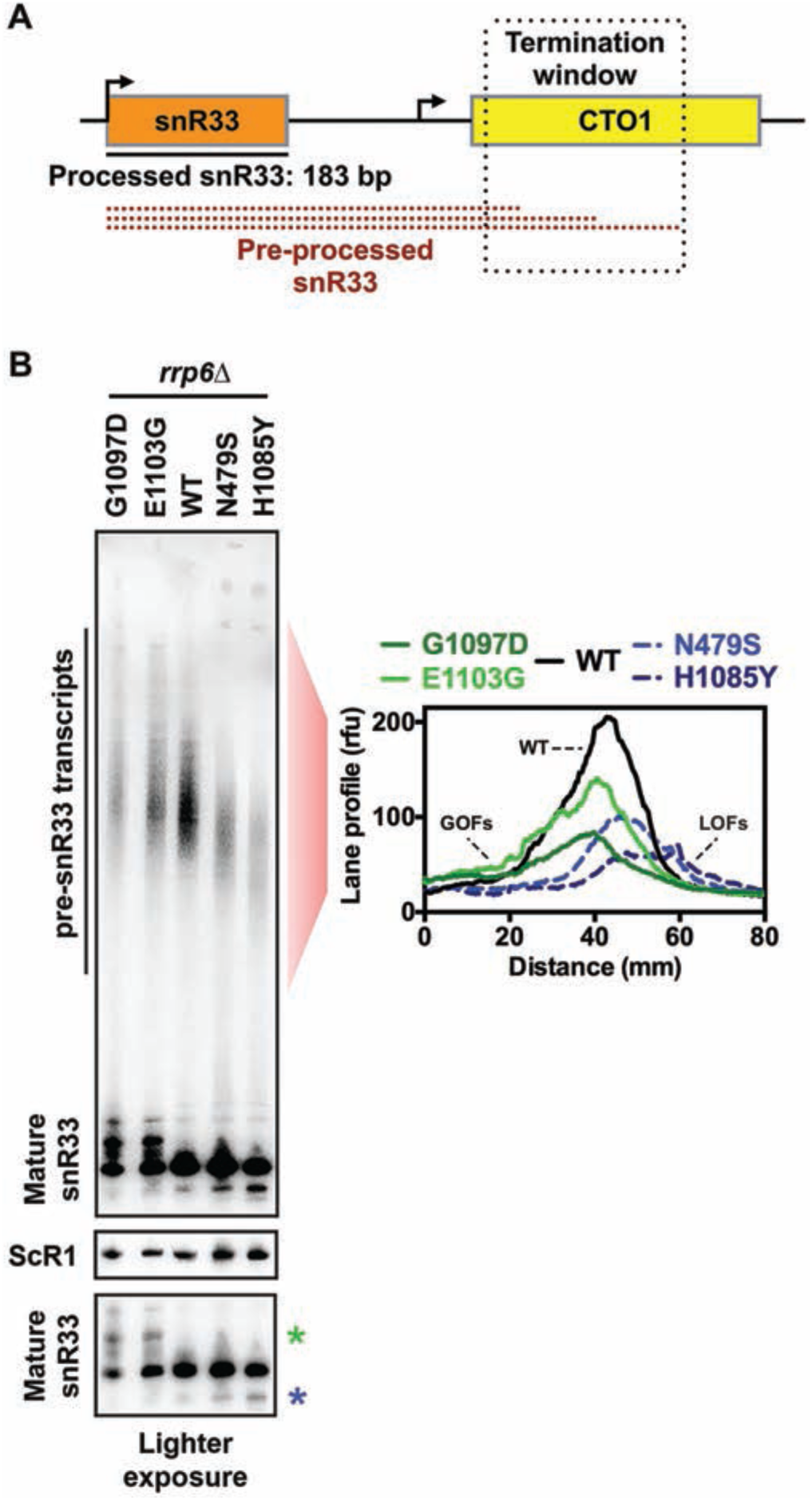
**Mapping of termination window for preprocessed *SNR33* in WT and Pol II catalytic mutants.** (A) Schematic of the snR33 termination window. (B) GOF and LOF mutants increase and decrease average length of pre-processed snR33, respectively. Northern blotting of *SNR33* for WT, slow (N479S and H1085Y) and fast (E1103G and G1097D) catalytic mutants containing deletion of the nuclear exosome subunit gene *RRP6.* Pol III transcript ScR1 was used as loading control. Average densitometric values of indicated lanes from three independent repeats are presented in the graph. A lighter exposure of the mature snR33 full-length transcript shows alternative longer (green asterisk) and shorter (blue asterisk) products for GOF and LOF mutants, respectively, that are proposed to generated by shifts in TSS usage.

### Glucose signal transduction defects in Pol II alleles may partly explain apparent slow elongation by ChIP

One contribution to apparent slower elongation of GOF mutants on the *GAL1*p*∷YLR454w* template could be delayed transcriptional shutoff. We reasoned that if glucose repression (or galactose starvation) were delayed in GOF mutants, it would add to apparent slow kinetics of Pol II exiting the gene. To this end, we investigated the kinetics of nuclear localization of Mig1p, which upon addition of glucose to medium is imported into the nucleus due to regulation by dephosphorylation, binds to target promoters of glucose-repressed genes and recruits co-repressors (64,65). As a proxy for early events in this cascade, we determined the kinetics of Mig1p nuclear localization by monitoring import of Mig1p-GFP upon glucose addition in WT or GOF catalytic mutants We observed signal for presumptive Mig1p-GFP localization to the nucleus saturate within ~2 min for WT cells, as previously reported (66). However, for at least one GOF mutant, the signal did not appear to reach maximum nuclear fluorescence within 2 min of glucose addition, thus we measured extended time points and analyzed the data from individual cells using nonlinear regression (Figure 7A and B). GOF G1097D showed both a delay in average time of Mig1p-GFP nuclear import and reduced average maximum nuclear fluorescence compared to WT, suggesting *in vivo* shutoff kinetics may be delayed due to deranged Mig1p signaling in this mutant at least (Figure 7B). Next, we measured extended time points of Mig1p translocation upon glucose repression over populations of individual cells for the GOF mutants analyzed in Figure 7A/B, while also including one LOF mutant, N479S. A noticeable subset of delayed responding cells was observed for GOF mutant G1097D, while higher fractions of non-responding cells were observed for both GOF mutants and the LOF mutant N479S compared to WT (Figure 7C and **Supplementary Figure 4**). Both GOF mutant G1097D and LOF mutant N479S showed lower average maximum nuclear Mig1p-GFP accumulation compared to WT, with the LOF mutant N479S showing an increase, then decrease, in Mig1p-GFP signal over time (**Supplementary Figure 4**). Altogether, these results indicate that Mig1p nuclear translocation in Pol II catalytic mutants can be aberrant, potentially affecting interpretation of transcription kinetics upon glucose shutoff. However, the strengths of observed apparent signaling defects do not appear to explain the large apparent fractions of retained mutant Pol II molecules at the 3′ end of *GAL1*p*∷YLR454w*.

**Figure 7.**
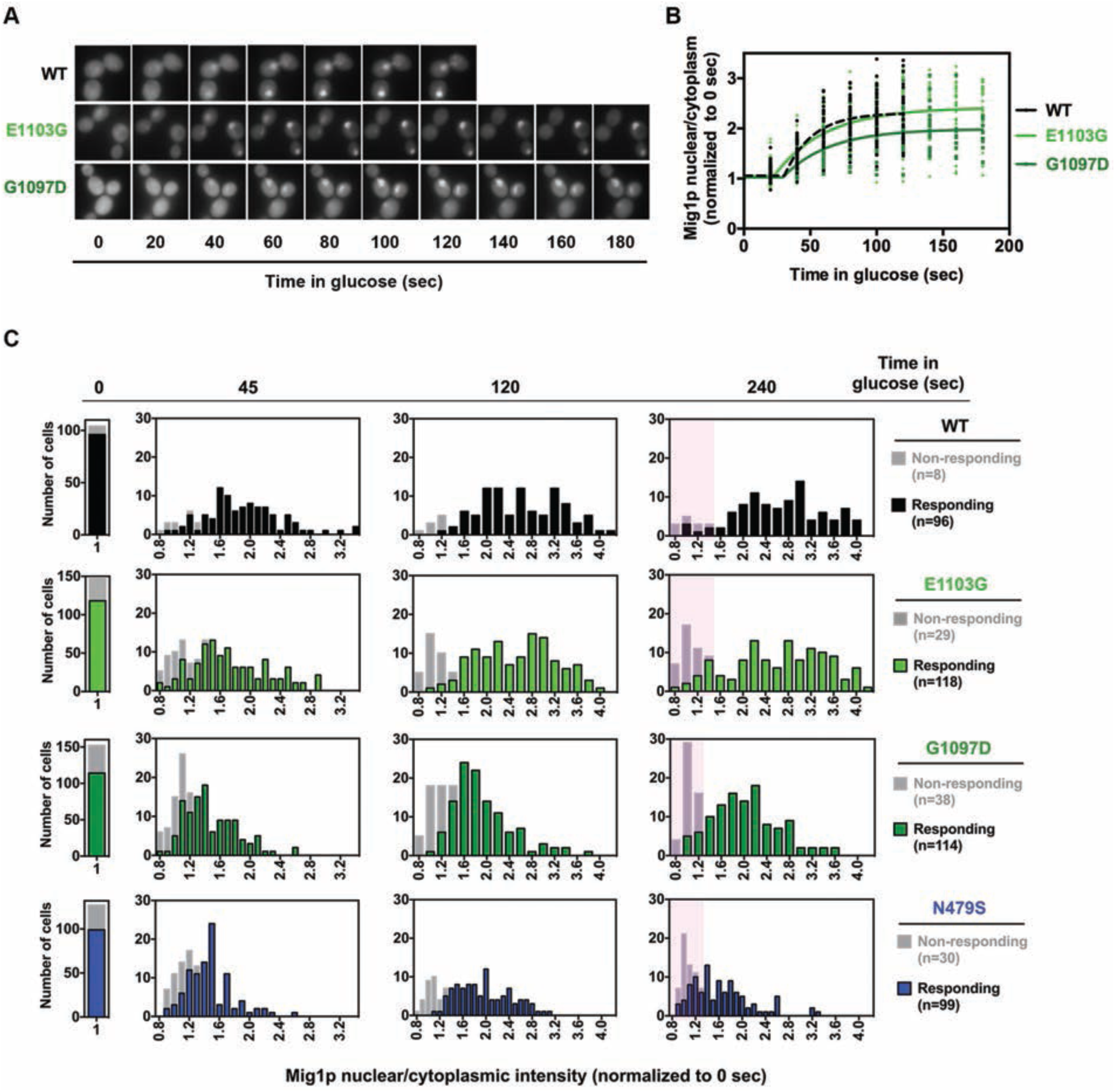
**Mig1 nuclear translocation is aberrant in Pol II catalytic mutants.** (A) Representative images of nuclear localization of Mig1p-GFP in WT and catalytic mutants (E1103G and G1097D) upon glucose (4% final) addition. Overnight grown cells were inoculated into fresh synthetic complete medium containing galactose (SC-2% Gal) and grown until mid-log phase at 30˚C. Pre-glucose repression samples were used as time 0 (t0), followed by replacing the medium with SC-2% Gal + 4% glucose to induce repression. (B) Mig1p nuclear localization kinetics is delayed in GOF mutant G1097D. Normalized (to pre-glucose treatment; t0) data from glucose-responding cells (n>35) plotted using nonlinear regression in GraphPad Prism. (C) Histograms of Mig1p nuclear localization show the distribution of nuclear Mig1p fluorescence (normalized to t0) intensity over indicated time points. Cells that do not show any traces of fluorescence accumulation are designated as non-responding cells (see methods for quantification details). Highlighted areas indicate population of cells that are either non-responding or responding cells with decreased Mig1p nuclear localization.

### Pol II catalytic mutants do not confer sensitization of cells to GTP starvation through hypothesized global elongation defects

To understand the relationship between altered Pol II activity and growth defects derived from putative global elongation defects, we turned to pharmacological reduction of a Pol II substrate, GTP. Nucleotide depleting drugs such as MPA or 6-AU have been interpreted as specifically exacerbating global elongation defects of Pol II or transcription factor mutants, and therefore have been widely used as pharmacological probes for transcription elongation *in vivo*. MPA depletes GTP through inhibition of the inosine monophosphate dehydrogenase (IMPDH) activities encoded by two MPA-sensitive IMPDH paralogs, *IMD3* and *IMD4.* WT cells are resistant to MPA because they are able to induce an MPA-resistant IMPDH-encoding paralog *IMD2* upon MPA-induced GTP starvation (Figure 8A). Interpretations of MPA sensitivity assume that WT and mutant strains have similar GTP starvation upon MPA treatment. We and others have noted that WT and MPA-sensitive mutants can experience different levels of GTP starvation due to different expression levels of the *IMD2* gene product, which provides MPA-resistant IMPDH activity as noted above (67–69). These differences in *IMD2* expression explain why increased catalytic activity Pol II GOF mutants are MPA sensitive – they do not express *IMD2*. In contrast, Pol II LOF mutants constitutively express *IMD2*, and do not show MPA sensitivity. We chose to evaluate how Pol II responds to GTP starvation and determine if Pol II transcription is a critical determinant for cellular sensitivity to GTP starvation. To do so, we rendered GTP starvation independent of *IMD2* expression by deleting *IMD2.* Deleting *IMD2* is predicted to normalize GTP starvation between WT and Pol II mutants, allowing us to examine the assertion that GTP depletion exacerbates putative Pol II elongation defects *in vivo* (Figure 8A). Furthermore, to enable an *in vivo* readout for *IMD2* expression, we replaced the *IMD2* ORF with *HIS3* (Figure 8A, lower panel).

**Figure 8.**
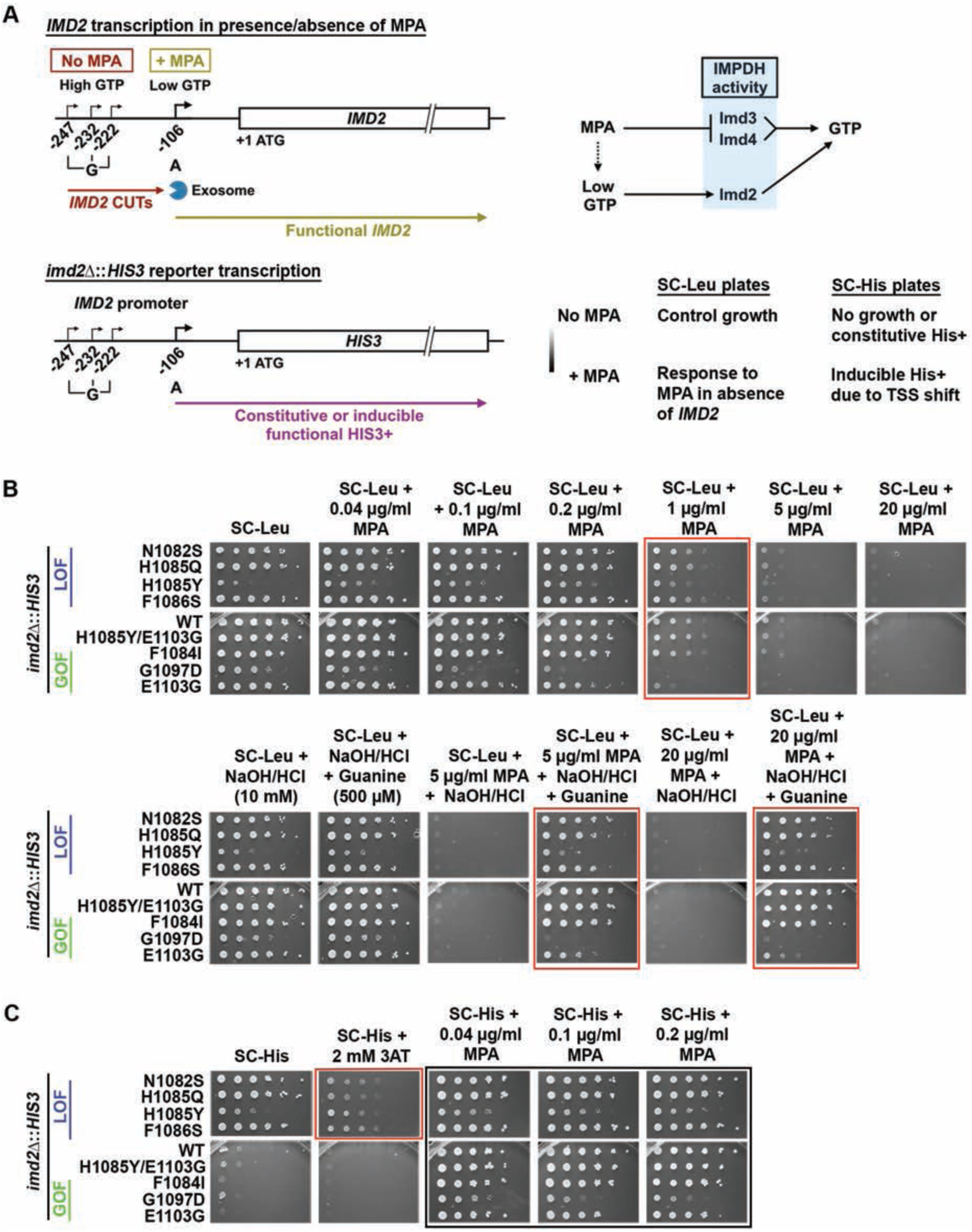
**Pol II catalytic mutants defective for *IMD2* transcription do not abolish GTP sensing.** (A) (top left) Schematic of *IMD2* gene transcription in presence and absence of MPA. In GTP-replete cells, *IMD2* is transcribed from upstream ‘G’ start sites producing non-functional CUTs, which are degraded by the RNA exosome. In the presence of MPA (GTP depletion), a downstream ‘A’ start site is utilized to produce functional *IMD2* transcripts. (top right) MPA depletes GTP by inhibition of IMPDH activity, present in yeast in two paralogous enzymes, Imd3 and Imd4. Upon GTP depletion, expression of an MPA-resistant IMPDH enzyme, Imd2 is induced. (bottom) Schematic of the *imd2*∆∷*HIS3* construct and expected phenotype upon MPA treatment in synthetic medium lacking leucine or histidine. All strains lack endogenous *IMD2* ORF, which is replaced with *HIS3* (*imd2*∆∷*HIS3*), hence rendering them highly sensitive to MPA treatment. (B) In absence of endogenous *IMD2*, Pol II catalytic mutants respond to GTP depletion similarly to WT. 10-fold serial dilutions of saturated cultures of Pol II catalytic mutants plated on synthetic medium lacking leucine for comparison of growth in various concentration of MPA treatment at 30˚C. (Upper panel) For the Pol II catalytic mutants’ response to MPA treatment, the most obvious comparison with WT can be made at 1 µg/ml MPA treatment (highlighted with red box). (Lower panel) Addition of guanine suppresses MPA sensitivity of Pol II mutants (highlighted with red box). (C) 10-fold serial dilutions of saturated cultures of Pol II catalytic mutants plated on synthetic medium lacking histidine for comparison of growth in various concentration of MPA treatment at 30˚C. Slow catalytic mutants constitutively use the downstream, productive ‘A’ site at the *IMD2* promoter, which produces a functional transcript (see text for details). Hence, His^+^ colonies are observed even in the absence of MPA (red highlighted box). All Pol II catalytic mutants show inducible His^+^ phenotype upon low level of MPA treatment (highlighted black boxes), indicating retention of GTP sensing ability.

In the absence of *IMD2*, otherwise wild type yeast strains become sensitive to MPA treatment as expected (Figure 8B) (67,68). If Pol II transcription were a major determinant of GTP starvation, it would be predicted that elongation-defective LOF Pol II alleles would become hypersensitive to MPA, while GOF increased catalytic activity mutants would show relative resistance. In contrast to these predictions, deletion of *IMD2* strongly blunted the Pol II allele-specific effects of MPA treatment on yeast cells (Figure 8B). We did observe mild MPA sensitivity for Pol II GOF alleles E1103G and G1097D. These results suggest that the major determinant for Pol II allele sensitivity to MPA is differential *IMD2* expression, and not necessarily Pol II elongation defects. As stated earlier, MPA effects are presumed to function through GTP starvation. This presumption can be tested by addition of guanine to the growth medium, which can suppress MPA-sensitivity by supporting an alternate route for GTP synthesis. In testing this presumption, we found that use of 10 mM NaOH to solubilize guanine in the medium on its own reduced effects of MPA on most of the Pol II mutants (**Supplemental Figure 5**). Notably, some of these Pol II mutants have shown allele-specific phenotypes upon change to pH of media (13). We determined that addition of equivalent HCl to neutralize NaOH effects allowed examination of guanine suppression of MPA sensitivity (Figure 8B). We observed that addition of guanine suppresses MPA sensitivity for most Pol II mutants except the GOF mutants E1103G and G1097D, which show mild or no suppression of MPA sensitivity by guanine, respectively. These results suggest that either the *IMD2*-independent MPA effects on GOF mutants are independent of GTP starvation, or these mutants have alterations in their GTP salvage synthesis pathways. We next determined if the ability to induce the *IMD2* promoter was maintained in Pol II mutants when GTP starvation conditions were normalized, using growth on medium lacking histidine to detect expression of *imd2∆∷HIS3* (Figure 8C). WT cells and GOF mutants were almost completely His^−^ in the absence of induction of the *IMD2* promoter (controlling *HIS3*), while LOF mutants were His^+^, indicative of constitutive expression of *imd2∆∷HIS3*. All strains showed induction of His^+^ phenotype in the presence of MPA (inducer of *IMD2* promoter), suggesting that productive transcription from the *IMD2* promoter was possible for all tested Pol II mutants (discussed further below).

A number of transcription factor mutants have been described as either MPA or 6-AU sensitive with these sensitivities interpreted as indicative of Pol II elongation defects. Such mutants include deletions in known elongation factor genes *dst1∆* (encodes TFIIS, (42,70)), *spt4∆* (subunit of Spt4/Spt5 DSIF, (71–73)), *paf1∆* (Paf1C complex member, (38,74)), *bur2∆* (P-TEFb subunit homolog, (72)), *pop2∆* (encodes Ccr4/NOT complex member, (42,75)) or deletions in genes encoding subunits of transcriptional coactivator complexes *spt3∆, sgf73∆* (SAGA, (38,42)) or *gal11∆* (Mediator, (42)). It has previously been demonstrated for some of these mutants, including *dst1*∆, *spt3*∆ and *paf1*∆, that *IMD2* induction is defective (38,42), indicating differential GTP starvation upon drug treatment for these strains relative to WT, just as for the Pol II mutants described above. We characterized these strains’ sensitivity to MPA treatment in presence or in the absence of endogenous *IMD2* (Figure 9). We observed that some of the mutants including *sgf73∆* and *pop2∆* were only slightly sensitive to MPA in presence of *IMD2* (Figure 9A, **top panels**). However, in the absence of *IMD2,* only *spt3∆* showed MPA hypersensitivity relative to WT, with *dst1∆* showing only slight sensitivity relative to WT. We next examined suppression of observed MPA sensitivities by guanine supplementation (Figure 9A, **bottom panels**). We observed that *spt3∆* was entirely suppressed by guanine, *dst1∆* mostly suppressed, and *spt4∆*, *paf1∆*, and *gal11∆* much less so. We hypothesize that *spt3∆* hypersensitivity relates to defects in *TPO1* expression, which is known to modulate MPA sensitivity (38). Taken together, these results are inconsistent with putative global Pol II transcription defects being a critical determinant for cell growth under GTP limitation. We further examined if *IMD2* promoter function in response to MPA were intact in the cohort of transcription factor mutants tested (Figure 9B). We observed a His^+^ phenotype consistent with constitutive expression of *imd2∆∷HIS3* in *spt4∆*. This phenotype suggests possible altered initiation in *spt4* mutants as observed in LOF Pol II mutants (Figure 8C). Furthermore, we observed that the His^+^ phenotype was inducible in the presence of MPA for all factor mutants except *spt3∆*. These results indicate that presumptive GTP sensing is maintained in most mutants, but that *spt3∆* cells have a distinct defect not observed for other factor mutants or Pol II mutants. This defect indicates an almost complete inability to induce the *IMD2* promoter∷*HIS3* reporter, although *spt3∆* is known to express *IMD2* upon MPA treatment at a very low level in presence of endogenous *IMD2* (42).

**Figure 9.**
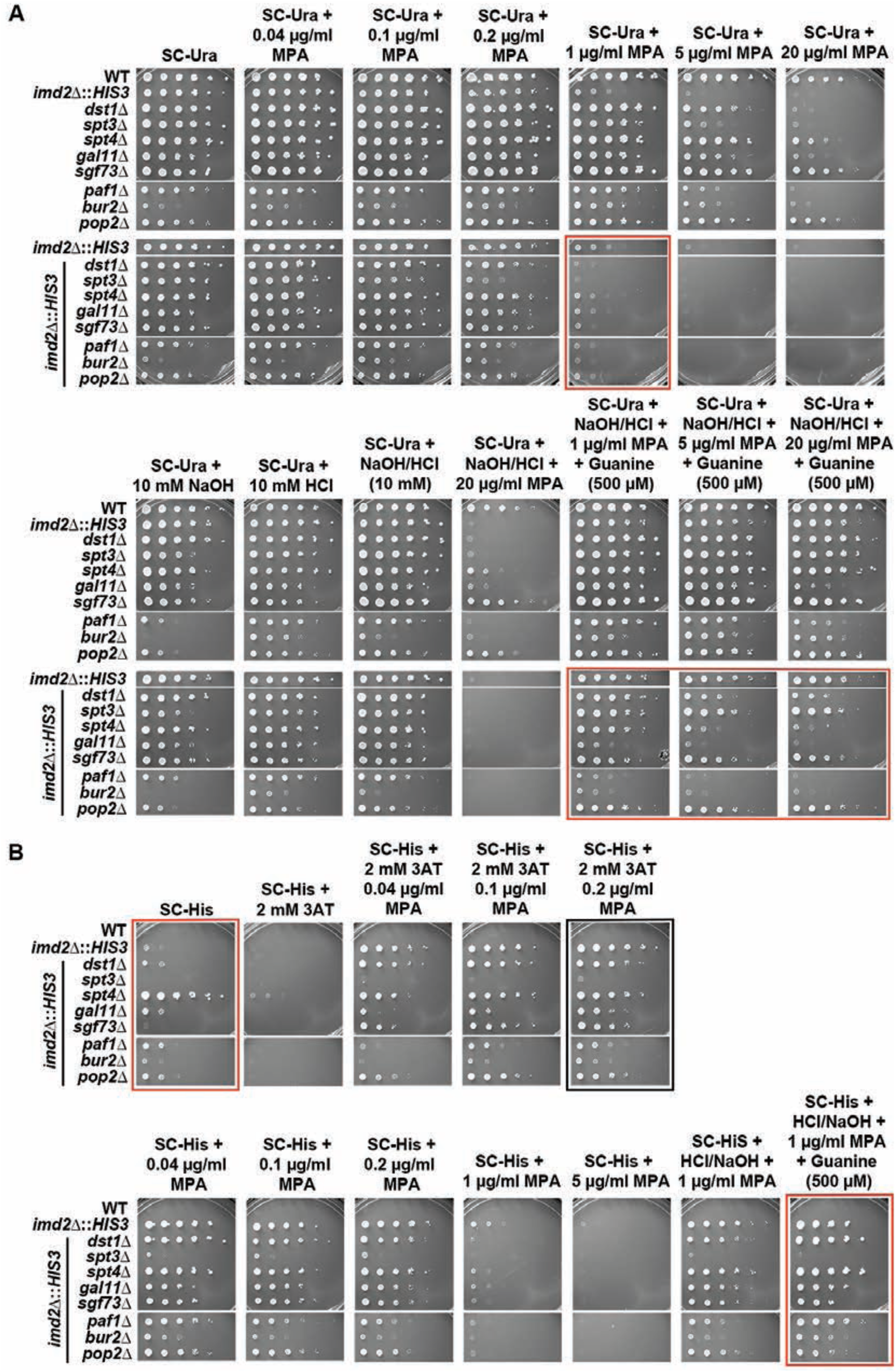
**Transcription related factor mutants respond similar way to WT upon MPA treatment in absence of endogenous *IMD2*.** (A) 10-fold serial dilutions of saturated cultures of indicated factor mutants plated on synthetic medium lacking uracil for comparison of growth in various concentration of MPA treatment at 30˚C. Upper and lower panels contain factor mutant strains with and without endogenous *IMD2* ORF (*imd2*∆∷*HIS3*), respectively. For convenience of comparison, spots of WT strain with *imd2*∆∷*HIS3* (second on the upper panel) is repeatedly presented next to factor mutants with *imd2*∆∷*HIS3* (top of the lower panel). In the absence of endogenous *IMD2*, most tested factor mutants show sensitivity to MPA at 1 µg/ml MPA treatment, which is comparable to WT sensitivity (upper panel highlighted red box). Addition of guanine suppresses MPA sensitivity for most of the factor mutants (highlighted red boxes, lower panel) (B) 10-fold serial dilutions of saturated cultures of indicated factor mutants plated on synthetic medium lacking histidine for comparison of growth in various concentration of MPA treatment at 30˚C. *spt4*∆ confers constitutive His^+^ phenotype (upper panel highlighted red box), while *spt3*∆ cannot show a His^+^ phenotype at the highest concentration of MPA used (upper panel highlighted black box). Addition of guanine suppresses MPA sensitivity of all factor mutants except *spt3*∆ (lower panel highlighted box).

### Pol II catalytic mutants do not abolish the response to GTP depletion but derange TSS usage at *IMD2* promoter

By removing the possibility of differential *IMD2* expression complicating the MPA response (using *imd2*∆), we showed above that many factor mutants or Pol II catalytic mutants have similar responses to MPA treatment. These results indicated Pol II transcription is not especially sensitive to presumptive GTP starvation *in vivo* relative to other pathways that rely on GTP (Figure 8 and 9). To better understand the mechanism of *IMD2* expression defects in the absence of differential GTP starvation, we further analyzed transcriptional responses at the *IMD2* promoter upon MPA treatment. Using the same *imd2∆*∷*HIS3* reporter construct used above, we analyzed the kinetics of TSS utilization upon addition of a concentration of MPA that induces a TSS shift at the *IMD2* promoter (Figure 10A). Pol II has been proposed to directly sense GTP levels through its active site, and through this sensing Pol II catalytic activity controls *IMD2* TSS usage (41). Under this model, it would be predicted that LOF Pol II mutants might show precocious downstream TSS usage; indeed, this behavior has previously been observed (4). Conversely, under this model, catalytically GOF Pol II mutants would show delayed TSS shifting kinetics due to their relative insensitivity to reduced GTP levels.

**Figure 10.**
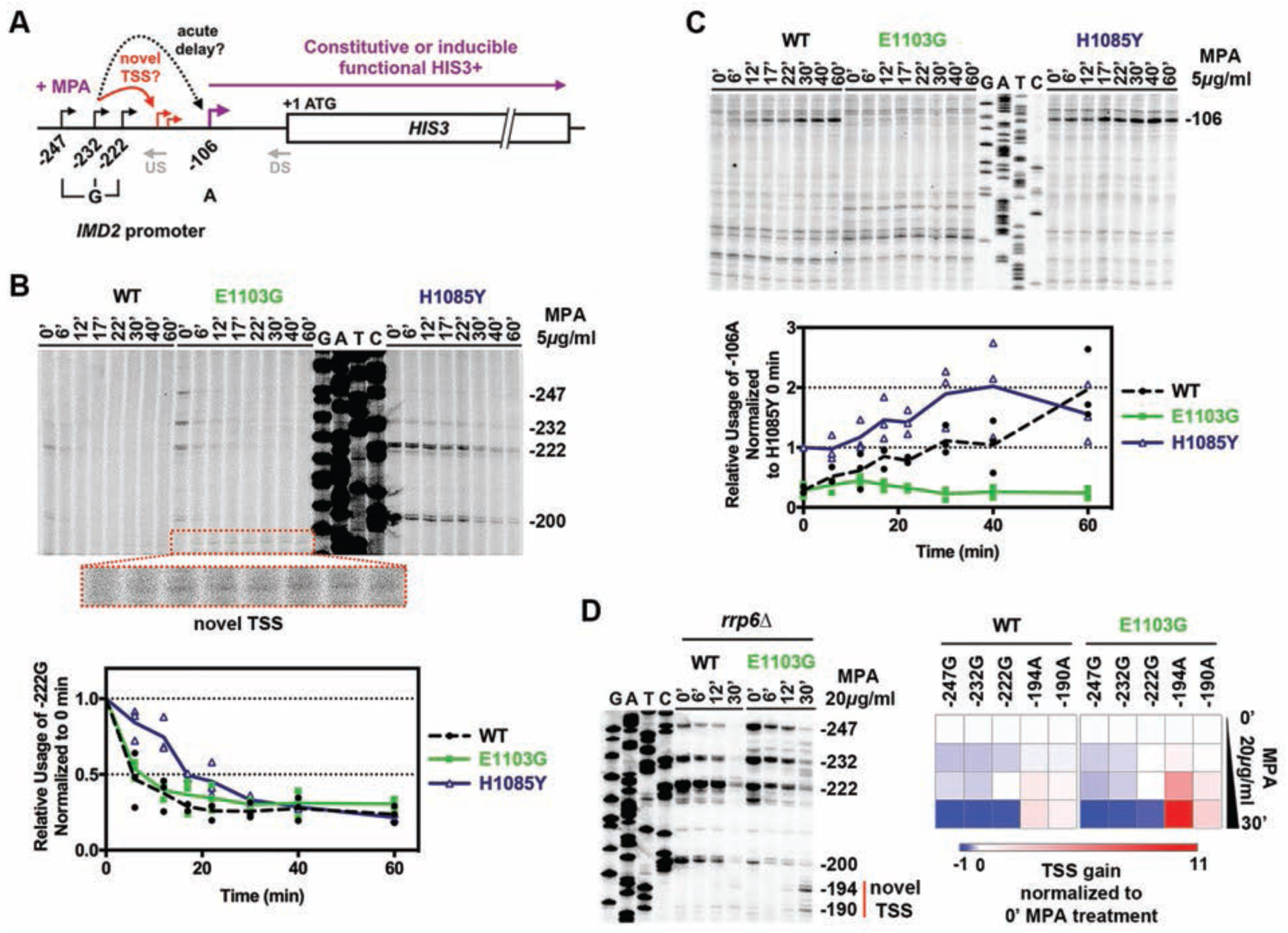
**Pol II catalytic mutants do not abolish response to GTP depletion.** (A) Schematic of the *imd2*∆∷*HIS3* construct used to determine WT and Pol II catalytic mutants’ response to nucleotide depletion in absence of endogenous *IMD2*. Architecture of the *IMD2* promoter indicating the upstream ‘G’ start sites that are used under normal GTP levels to produce non-functional *IMD2* cryptic unstable transcripts (CUTs). Downstream -106 ‘A’ is used upon presumptive GTP-depletion in presence of MPA, producing a functional transcript that confers MPA resistance if *IMD2* ORF is present. Slow catalytic mutants constitutively show use of both upstream and downstream start sites. Gray “US” (Upstream) and “DS” (Downstream) indicate positions of primers used for primer extension experiments. Two models for MPA-sensitivity of Pol II mutants are shown: TSS shift to inappropriate novel TSS, or acute delay in shift to appropriate TSS. (B) Both GOF and LOF mutants lose upstream ‘G’ TSSs upon MPA treatment. Time courses showing the usage of upstream ‘G’ sites upon MPA treatment (US primer used for PE). Cells grown to mid-log phase in synthetic complete media at 30˚C were treated with MPA (5 µg/mL final concentration) and RNA was isolated at indicated time points. Pre-treatment sample was used for time 0. Graph shows the relative usage of -222G site for WT and mutants upon MPA treatment normalized to 0 min (no-treatment). (C) E1103G is unable to shift TSS usage to downstream functional ‘A’ site upon GTP depletion. Same RNAs from Figure 8B used for PE experiment with DS primer. Graph shows the relative gain of -106A site for WT and mutants upon MPA treatment normalized to H1085Y 0 min (no-treatment), which is used constitutively. (D) Fast catalytic mutants gradually lose upstream ‘G’ sites and shift TSS usage to novel ‘A’ sites (putatively -194A and -190A) in response to nucleotide depletion. Usage of these sites is likely produce non-functional CUTs, shown to be stabilized here by the deletion of exosome subunit gene *RRP6*. Heat map represents the average loss/gain of indicated TSS upon MPA treatment obtained from three independent repeats (see **Supplementary Fig 6**).

We observed that both LOF (H1085Y) and GOF (E1103G) mutants gradually lose upstream ‘G’ sites upon MPA treatment (Figure 10B), indicating retention of ability to respond to GTP-depletion. As expected, WT cells rapidly lose upstream ‘G’ TSSs and subsequently gain the downstream functional ‘A’ TSS (Figure 10C). H1085Y constitutively uses the downstream functional ‘A’ TSS and shows increased usage of this site upon MPA treatment, while showing decreased kinetics of loss of upstream TSSs. This result is unexpected from the “defective GTP sensing” model where kinetics of upstream TSS loss are predicted to be faster than WT, not slower for Pol II LOF mutants (Fig 10 B and C). E1103G shows reduced kinetics of loss of upstream site, consistent with defective GTP sensing, but shifts TSS usage to novel TSSs predicted to be nonfunctional due to their position upstream of the *IMD2* Nab3/Nrd1-dependent terminator (Figure 10B and C). These novel TSSs are upstream of the normal −106 A *IMD2* TSS, and their appearance in GOF E1103G are consistent with E1103G shifting TSSs upstream at other genes (4). If terminated by the Nab/Nrd pathway, putative non-functional novel TSSs should produce cryptic unstable transcripts (CUTs) that are degraded by the nuclear exosome (76,77). Such *IMD2* CUTs have previously been observed in *sen1*, *nab3* and *rpb11* mutants (41). Indeed, deletion of the exosome subunit Rrp6 also stabilized CUTs derived from the novel TSSs we observe, with significant utilization of these TSSs in E1103G compared to WT (Figure 10D and **Supplementary Figure 6A-B**). Stabilization of these CUTs in *rrp6*∆ strains did not provide MPA resistance for MPA sensitive Pol II GOF mutants, as these transcripts do not likely produce functional *IMD2* transcript required for resistance even if stabilized (**Supplementary Figure 6C**). We note that although E1103G generally appears unable to utilize downstream functional ‘A’ TSS upon MPA treatment, enough functional transcript was made to produce a His^+^ phenotype in the presence of *imd2∆∷HIS3* at low concentrations of MPA treatment (Figure 9B). Our results here provide an initiation-based mechanism for the MPA sensitivity of Pol II GOF mutants.

## Discussion

Examination of Pol II catalytic mutants by our lab and others has shown Pol II activity-sensitive aspects of transcription initiation, elongation, co-transcriptional processes, and termination (4,7,8,15,19,20). Our current study utilizes a set of yeast Pol II catalytic mutants to probe broad aspects of gene expression *in vivo*. We show that Pol II catalytic defects lead to a decrease in overall Pol II occupancy with reduction in 3’ end occupancy on a galactose inducible reporter gene, *GAL1*p∷*YLR454w* (Figure 1). Further, using a constitutively expressed *TEF1*p∷*YLR454w* reporter we find that growth on different carbon sources may alter Pol II occupancy profiles (Figure 1). Previously, several observations suggested alterations of Pol II activity and gene expression due to external perturbations such as carbon source or temperature (54,55). Likewise, environmental stress has also been shown to affect Pol II activity and gene expression (78). Notably, it has also been shown that RAS/PKA signaling pathway, which controls aspects of glucose signaling, can target proteins associated with general transcription machinery, can putatively regulate elongating Pol II by targeting Spt5/4 (79,80), and shows interactions with Nab3/Nrd1 termination factors (81). In addition to differential effects of carbon source on steady state Pol II occupancy, we also observe impaired galactose induction and glucose repression kinetics in both LOF and GOF mutants (Figure 2 and 7). Interestingly, both classes of catalytic mutants show impaired growth at 37˚C on galactose likely arising from the sum of a number of distinct individual defects in initiation, elongation, and termination (**Supplementary Figure 1B**). Taken together, these results support growth condition-dependent modulation Pol II elongation *in vivo*.

Pol II mutant *rpb1* E1103G, which was reported previously to be fast for elongation *in vivo* relative to WT (19), appears slower than WT in our study. *In vitro* biochemical studies, including direct observations of individual polymerases at the single molecule level, have repeatedly shown faster elongation rate for E1103G compared to WT (4,6,12,14). Several lines of evidence suggest E1103G *in vitro* GOF activity manifests itself *in vivo* through transcriptional effects distinct from LOF mutants. First, primer extension analysis shows that E1103G shifts TSS usage upstream at a number of promoters, consistent with a model that increase in catalytic activity increases initiation probability during Pol II promoter scanning, resulting in observed upstream TSS shifts (4,7). Second, E1103G and other Pol II GOF mutants show allele-specific genetic interactions with a number of factors, suggesting their defects are distinct from LOF mutants (7). For example, the synthetic sick genetic interaction of E1103G with the RNA processing factor alleles *rat1-1* and *xrn1∆*/*rat1-1* is consistent with exacerbation of termination defects through faster elongation (18–20,60), and in contrast to the suppression of *rat1-1* and *rat1-1/xrn1∆* by LOF Pol II alleles. Indeed, delayed termination for GOF E1103G was evident through observation of increased length of pre-processed snR33 (Figure 6). We further show that an even stronger GOF mutant, G1097D, shows a corresponding increase in pre-processed snR33 length relative to E1103G or WT. While increased length of pre-processed snR33 is consistent with increased elongation rate in GOF mutants, defective termination due to altered probability of a Pol II conformational change should not be ruled out as possible mechanism (21). Finally, a recent study demonstrated that an E1103G strain displays a shift in co-transcriptional splicing towards downstream positions, consistent Pol II traveling further downstream prior to splicing, though delayed splicing kinetics cannot be ruled out (16). Altogether, these data are consistent with increased elongation kinetics in GOF catalytic mutants *in vivo* under a parsimonious view, though they do not explain the delayed Pol II runoff from *YLR454w* observed here. We speculate that template, context (such as distance from promoter, position within gene, or gene expression level), or condition-specific effects may contribute to either slow elongation or template retention in fast Pol II catalytic mutants in some genomic locations.

Our results suggest that the commonly used glucose shutoff assay for determining *in vivo* elongation rate may be confounded by previously unappreciated limitations and biological complexity. We also identify a possible artifact of prior experimental design that may have contributed to discrepant results reported for GOF mutant E1103G. First, we have shown that galactose starvation induces as fast a transcriptional shutoff at *GAL1*p∷*YLR454w* as glucose addition (Figure 5F). Second, this galactose-depleted transcriptional shutoff affects Pol II E1103G differently than WT. During the wash, E1103G Pol II runs off both the 5’ and 3’ ends of the reporter faster than WT (Figure 5F-G). 5’ runoff is consistent with E1103G elongating faster than WT early in the time course, or E1103G being more sensitive to galactose starvation. The increased runoff kinetics from the 3’ end of the reporter in E1103G is perplexing. Furthermore, after these early effects observed for E1103G relative to WT under galactose starvation, E1103G Pol II shows a delay in running off the template at longer time points regardless of mechanism of shutoff, just as purported LOF Pol II alleles do under glucose inhibition. From these results we can only conclude that Pol II could be differentially affected while on the 3’ end of the gene relative to the middle under different growth conditions. Further, we identified slow Mig1p nuclear translocation upon glucose exposure as a possible confounding variable for use of *GAL1*p∷*YLR454w* to determine *in vivo* elongation rate. There may be additional untested steps in inhibition specific to the *GAL* system that are defective, or defects specific to the long *YLR454w* template for Pol II GOF strains. Regardless, careful consideration of possible variables in assay behavior is urged given the results we present.

In order to exploit the existing transcription reporter systems to probe transcription mechanisms carefully, we need a detailed understanding of each phenotypic system, such as our analyses to dissect the Spt^-^ phenotype of Pol II GOF alleles (82). Here we investigated Pol II catalytic mutants and several transcription factors mutants’ response to the nucleotide-depleting drug MPA under conditions where differential *IMD2* expression was obviated. Most tested Pol II mutants and many transcription factor mutants behave similarly to WT upon MPA treatment of *imd2∆* strains (Figure 8 and 9). Our results also suggest *IMD2*-independent mechanisms of MPA sensitivity, which have been observed in previous large-scale deletion screens for MPA or 6-AU sensitivity, and have identified Pol II mutants and transcription factor mutants that do not affect *IMD2* transcription (38,42). Our previous analyses identified a correlation between MPA sensitivity of Pol II GOF alleles and upstream shifts in TSS usage at *ADH1* (4,7). As *IMD2* regulation proceeds by initiation shifting from upstream non-productive TSSs to a downstream productive one, it is conceivable that initiation defects underlie Pol II mutant MPA sensitivity. However, in the presence of endogenous *IMD2* it is difficult to determine mechanisms of MPA sensitivity of GOF mutants, as they would be differentially starved for GTP due to differential expression of functional *IMD2* upon MPA treatment. One model for MPA sensitivity based on differential sensitivity to reduced GTP is that increased catalysis rate of GOF alleles might buffer reduction in substrate (GTP) levels. This could lead to delay in sensing low GTP and switching *IMD2* upstream GTP-initiated TSSs to downstream ATP-initiated TSSs, causing acute GTP starvation possibly beyond a critical threshold for growth/viability. Our results disfavor this model. Here we present evidence that MPA sensitivity of the Pol II GOF allele E1103G correlates with usage of novel TSSs that are intermediately positioned between the known productive -106 A TSS at *IMD2* and upstream non-functional starts. We conclude that *IMD2* defects in Pol II GOF E1103G are likely to derive from initiation defects (Figure 10).

Our Pol II catalytic mutants and pre-mRNA processing factor show allele-specific genetic interactions (Figure 3). Synthetic sick interactions of GOF with *xrn1*∆ and stabilization of the *GAL1*p∷*YLR454w* transcript in the *xrn1*∆/*rat1-1* double mutant background is suggestive of a 5’-end processing defect in GOF mutants, possibly a capping defect. A defect in capping would be predicted to expose mRNA to the action of 5’ exonucleases Xrn1 and Rat1. Capping of nascent transcripts occurs co-transcriptionally with Pol II elongation proposed to be coupled to successful capping, though a direct “capping checkpoint” has yet to be shown (83,84). Potential sensitivity of capping to Pol II GOF mutants is suggestive of either a defective checkpoint in Pol II mutant strains or the absence of one. We also observe correlations of reporter transcript half-life with expression levels in Pol II mutants, and with Pol II mutants’ growth defects (Figure 4). Observed correlations between mRNA decay rate with both expression and growth rate is consistent with recent findings (27)(http://biorxiv.org/content/early/2016/03/19/044735). ‘Feedback’ of gene expression control between the abundance of mRNA and mRNA decay rates is proposed to occur in response to defects in transcription synthesis (24,25). Alternatively, as growth efficiency is connected to overall translational demand, mRNA decay ‘feedback’ may occur upon alteration of overall translation rate, as most mRNA decay happens co-translationally (85–87). Given that Pol II mutants show reduced apparent initiation based on overall Pol II occupancy and reduced growth rates, it is difficult to deconvolute primary and secondary global effects. Although we looked at both phenomena using a single reporter transcript, mutants described here could be used further to probe these proposed mechanisms in large scale.

Here we extensively characterize a set of Pol II catalysis mutants for *in vivo* consequences. We show that altered Pol II catalysis affects Pol II occupancy, putative elongation, and reporter gene expression and decay rate *in vivo*. Notably, we interrogate two widely used elongation reporter systems, raising caveats about their use and interpretation. For use of nucleotide depleting drugs MPA or 6-AU, we constructed and tested a useful novel reporter system (*imd2*∆*∷HIS3*), which can be further utilized to characterize or screen for new mutants that shift TSS usage downstream, leading to constitutively expression of *imd2∆∷HIS3*. Development of approaches allowing more direct determination of *in vivo* elongation rate will bypass issues identified here. Recent advances in high-resolution microscopy have enabled real-time observation of all transcription phases on endogenous genes using fluorescently labeled proteins that bind to nascent transcript (88,89). This approach could be to address how any number of variables might modulate elongation such as template sequence, RNA secondary structure *etc*, and likely represents the next steps toward understanding transcription elongation and co-transcriptional processes *in vivo*.

## Acknowledgments

We thank Mary Bryk for providing bead beater for yeast cell breakage and UV cross-linker for blotting. We are grateful to Jennifer Herman and Lanying Zeng for providing microscope time and training. We thank Ry Young/Center for Phage Technology for use of Tecan plate reader. We thank Kaplan lab members for critical reading of the manuscript and helpful comments. This work was supported by grant R01GM097260 from the National Institutes of Health, National Institute for General Medical Sciences to CDK, and grant A-1763 from the Welch Foundation to CDK.

## References

1. Kaplan, C.D. (2013) Basic mechanisms of RNA polymerase II activity and alteration of gene expression in Saccharomyces cerevisiae. Biochim Biophys Acta, 1829, 39-54.

2. Svetlov, V. and Nudler, E. (2013) Basic mechanism of transcription by RNA polymerase II. Biochim Biophys Acta, 1829, 20-28.

3. Bentley, D.L. (2014) Coupling mRNA processing with transcription in time and space. Nat Rev Genet, 15, 163-175.

4. Kaplan, C.D., Jin, H., Zhang, I.L. and Belyanin, A. (2012) Dissection of Pol II trigger loop function and Pol II activity-dependent control of start site selection in vivo. PLoS Genet, 8, e1002627.

5. Kaplan, C.D., Larsson, K.M. and Kornberg, R.D. (2008) The RNA polymerase II trigger loop functions in substrate selection and is directly targeted by alpha-amanitin. Mol Cell, 30, 547-556.

6. Malagon, F., Kireeva, M.L., Shafer, B.K., Lubkowska, L., Kashlev, M. and Strathern, J.N. (2006) Mutations in the Saccharomyces cerevisiae RPB1 gene conferring hypersensitivity to 6-azauracil. Genetics, 172, 2201-2209.

7. Braberg, H., Jin, H., Moehle, E.A., Chan, Y.A., Wang, S., Shales, M., Benschop, J.J., Morris, J.H., Qiu, C., Hu, F. et al. (2013) From structure to systems: high-resolution, quantitative genetic analysis of RNA polymerase II. Cell, 154, 775-788.

8. Jin, H. and Kaplan, C.D. (2014) Relationships of RNA polymerase II genetic interactors to transcription start site usage defects and growth in Saccharomyces cerevisiae. G3 (Bethesda), 5, 21-33.

9. Qiu, C., Erinne, O.C., Dave, J.M., Cui, P., Jin, H., Muthukrishnan, N., Tang, L.K., Babu, S.G., Lam, K.C., Vandeventer, P.J. et al. (2016) High-Resolution Phenotypic Landscape of the RNA Polymerase II Trigger Loop. PLoS Genet, 12, e1006321.

10. Wang, D., Bushnell, D.A., Westover, K.D., Kaplan, C.D. and Kornberg, R.D. (2006) Structural basis of transcription: role of the trigger loop in substrate specificity and catalysis. Cell, 127, 941-954.

11. Kireeva, M.L., Nedialkov, Y.A., Cremona, G.H., Purtov, Y.A., Lubkowska, L., Malagon, F., Burton, Z.F., Strathern, J.N. and Kashlev, M. (2008) Transient reversal of RNA polymerase II active site closing controls fidelity of transcription elongation. Mol Cell, 30, 557-566.

12. Larson, M.H., Zhou, J., Kaplan, C.D., Palangat, M., Kornberg, R.D., Landick, R. and Block, S.M. (2012) Trigger loop dynamics mediate the balance between the transcriptional fidelity and speed of RNA polymerase II. Proc Natl Acad Sci U S A, 109, 6555-6560.

13. Cabart, P., Jin, H., Li, L. and Kaplan, C.D. (2014) Activation and reactivation of the RNA polymerase II trigger loop for intrinsic RNA cleavage and catalysis. Transcription, 5, e28869.

14. Dangkulwanich, M., Ishibashi, T., Liu, S., Kireeva, M.L., Lubkowska, L., Kashlev, M. and Bustamante, C.J. (2013) Complete dissection of transcription elongation reveals slow translocation of RNA polymerase II in a linear ratchet mechanism. Elife, 2, e00971.

15. Fong, N., Kim, H., Zhou, Y., Ji, X., Qiu, J., Saldi, T., Diener, K., Jones, K., Fu, X.D. and Bentley, D.L. (2014) Pre-mRNA splicing is facilitated by an optimal RNA polymerase II elongation rate. Genes Dev, 28, 2663-2676.

16. Carrillo Oesterreich, F., Herzel, L., Straube, K., Hujer, K., Howard, J. and Neugebauer, K.M. (2016) Splicing of Nascent RNA Coincides with Intron Exit from RNA Polymerase II. Cell, 165, 372-381.

17. de la Mata, M., Alonso, C.R., Kadener, S., Fededa, J.P., Blaustein, M., Pelisch, F., Cramer, P., Bentley, D. and Kornblihtt, A.R. (2003) A slow RNA polymerase II affects alternative splicing in vivo. Mol Cell, 12, 525-532.

18. Jimeno-Gonzalez, S., Haaning, L.L., Malagon, F. and Jensen, T.H. (2010) The yeast 5’-3’ exonuclease Rat1p functions during transcription elongation by RNA polymerase II. Mol Cell, 37, 580-587.

19. Hazelbaker, D.Z., Marquardt, S., Wlotzka, W. and Buratowski, S. (2013) Kinetic competition between RNA Polymerase II and Sen1-dependent transcription termination. Mol Cell, 49, 55-66.

20. Fong, N., Brannan, K., Erickson, B., Kim, H., Cortazar, M.A., Sheridan, R.M., Nguyen, T., Karp, S. and Bentley, D.L. (2015) Effects of Transcription Elongation Rate and Xrn2 Exonuclease Activity on RNA Polymerase II Termination Suggest Widespread Kinetic Competition. Mol Cell, 60, 256-267.

21. Zhang, H., Rigo, F. and Martinson, H.G. (2015) Poly(A) Signal-Dependent Transcription Termination Occurs through a Conformational Change Mechanism that Does Not Require Cleavage at the Poly(A) Site. Mol Cell, 59, 437-448.

22. Braun, K.A., Vaga, S., Dombek, K.M., Fang, F., Palmisano, S., Aebersold, R. and Young, E.T. (2014) Phosphoproteomic analysis identifies proteins involved in transcription-coupled mRNA decay as targets of Snf1 signaling. Sci Signal, 7, ra64.

23. Braun, K.A. and Young, E.T. (2014) Coupling mRNA synthesis and decay. Mol Cell Biol, 34, 4078-4087.

24. Haimovich, G., Medina, D.A., Causse, S.Z., Garber, M., Millan-Zambrano, G., Barkai, O., Chavez, S., Perez-Ortin, J.E., Darzacq, X. and Choder, M. (2013) Gene expression is circular: factors for mRNA degradation also foster mRNA synthesis. Cell, 153, 1000-1011.

25. Sun, M., Schwalb, B., Pirkl, N., Maier, K.C., Schenk, A., Failmezger, H., Tresch, A. and Cramer, P. (2013) Global analysis of eukaryotic mRNA degradation reveals Xrn1-dependent buffering of transcript levels. Mol Cell, 52, 52-62.

26. Haimovich, G., Choder, M., Singer, R.H. and Trcek, T. (2013) The fate of the messenger is predetermined: a new model for regulation of gene expression. Biochim Biophys Acta, 1829, 643-653.

27. Garcia-Martinez, J., Delgado-Ramos, L., Ayala, G., Pelechano, V., Medina, D.A., Carrasco, F., Gonzalez, R., Andres-Leon, E., Steinmetz, L., Warringer, J. et al. (2016) The cellular growth rate controls overall mRNA turnover, and modulates either transcription or degradation rates of particular gene regulons. Nucleic Acids Res, 44, 3643-3658.

28. Neymotin, B., Ettorre, V. and Gresham, D. (2016) Multiple Transcript Properties Related to Translation Affect mRNA Degradation Rates in Saccharomyces cerevisiae. G3 (Bethesda), 6, 3475-3483.

29. Mason, P.B. and Struhl, K. (2005) Distinction and relationship between elongation rate and processivity of RNA polymerase II in vivo. Mol Cell, 17, 831-840.

30. Hirayoshi, K. and Lis, J.T. (1999) Nuclear run-on assays: assessing transcription by measuring density of engaged RNA polymerases. Methods Enzymol, 304, 351-362.

31. Garcia-Martinez, J., Aranda, A. and Perez-Ortin, J.E. (2004) Genomic run-on evaluates transcription rates for all yeast genes and identifies gene regulatory mechanisms. Mol Cell, 15, 303-313.

32. Morillo-Huesca, M., Vanti, M. and Chavez, S. (2006) A simple in vivo assay for measuring the efficiency of gene length-dependent processes in yeast mRNA biogenesis. FEBS J, 273, 756-769.

33. Ardehali, M.B. and Lis, J.T. (2009) Tracking rates of transcription and splicing in vivo. Nat Struct Mol Biol, 16, 1123-1124.

34. Archambault, J., Lacroute, F., Ruet, A. and Friesen, J.D. (1992) Genetic interaction between transcription elongation factor TFIIS and RNA polymerase II. Mol Cell Biol, 12, 4142-4152.

35. Sweeney, M.J. (1977) Mycophenolic acid and its mechanism of action in cancer and psoriasis. Jpn J Antibiot, 30 Suppl, 85-92.

36. Powell, W. and Reines, D. (1996) Mutations in the second largest subunit of RNA polymerase II cause 6-azauracil sensitivity in yeast and increased transcriptional arrest in vitro. J Biol Chem, 271, 6866-6873.

37. Reines, D. (2003) Use of RNA yeast polymerase II mutants in studying transcription elongation. Methods Enzymol, 371, 284-292.

38. Desmoucelles, C., Pinson, B., Saint-Marc, C. and Daignan-Fornier, B. (2002) Screening the yeast “disruptome” for mutants affecting resistance to the immunosuppressive drug, mycophenolic acid. J Biol Chem, 277, 27036-27044.

39. Shaw, R.J. and Reines, D. (2000) Saccharomyces cerevisiae transcription elongation mutants are defective in PUR5 induction in response to nucleotide depletion. Mol Cell Biol, 20, 7427-7437.

40. Jenks, M.H., O’Rourke, T.W. and Reines, D. (2008) Properties of an intergenic terminator and start site switch that regulate IMD2 transcription in yeast. Mol Cell Biol, 28, 3883-3893.

41. Kuehner, J.N. and Brow, D.A. (2008) Regulation of a eukaryotic gene by GTP-dependent start site selection and transcription attenuation. Mol Cell, 31, 201-211.

42. Riles, L., Shaw, R.J., Johnston, M. and Reines, D. (2004) Large-scale screening of yeast mutants for sensitivity to the IMP dehydrogenase inhibitor 6-azauracil. Yeast, 21, 241-248.

43. Amberg, D.C., Burke, D.J. and Strathern, J.N. (2005) Methods in Yeast Genetics: A Cold Spring Harbor Laboratory Course Manual, 2005 Edition (Cold Spring).

44. Lenstra, T.L., Benschop, J.J., Kim, T., Schulze, J.M., Brabers, N.A., Margaritis, T., van de Pasch, L.A., van Heesch, S.A., Brok, M.O., Groot Koerkamp, M.J. et al. (2011) The specificity and topology of chromatin interaction pathways in yeast. Mol Cell, 42, 536-549.

45. Kaplan, C.D., Holland, M.J. and Winston, F. (2005) Interaction between transcription elongation factors and mRNA 3’-end formation at the Saccharomyces cerevisiae GAL10-GAL7 locus. J Biol Chem, 280, 913-922.

46. Livak, K.J. and Schmittgen, T.D. (2001) Analysis of relative gene expression data using real-time quantitative PCR and the 2(-Delta Delta C(T)) Method. Methods, 25, 402-408.

47. Schmitt, M.E., Brown, T.A. and Trumpower, B.L. (1990) A rapid and simple method for preparation of RNA from Saccharomyces cerevisiae. Nucleic Acids Res, 18, 3091-3092.

48. Marquardt, S., Hazelbaker, D.Z. and Buratowski, S. (2011) Distinct RNA degradation pathways and 3’ extensions of yeast non-coding RNA species. Transcription, 2, 145-154.

49. Ranish, J.A. and Hahn, S. (1991) The yeast general transcription factor TFIIA is composed of two polypeptide subunits. J Biol Chem, 266, 19320-19327.

50. Miermont, A., Waharte, F., Hu, S., McClean, M.N., Bottani, S., Leon, S. and Hersen, P. (2013) Severe osmotic compression triggers a slowdown of intracellular signaling, which can be explained by molecular crowding. Proc Natl Acad Sci U S A, 110, 5725-5730.

51. Danko, C.G., Hah, N., Luo, X., Martins, A.L., Core, L., Lis, J.T., Siepel, A. and Kraus, W.L. (2013) Signaling pathways differentially affect RNA polymerase II initiation, pausing, and elongation rate in cells. Mol Cell, 50, 212-222.

52. Jonkers, I., Kwak, H. and Lis, J.T. (2014) Genome-wide dynamics of Pol II elongation and its interplay with promoter proximal pausing, chromatin, and exons. Elife, 3, e02407.

53. Veloso, A., Kirkconnell, K.S., Magnuson, B., Biewen, B., Paulsen, M.T., Wilson, T.E. and Ljungman, M. (2014) Rate of elongation by RNA polymerase II is associated with specific gene features and epigenetic modifications. Genome Res, 24, 896-905.

54. Miguel, A., Monton, F., Li, T., Gomez-Herreros, F., Chavez, S., Alepuz, P. and Perez-Ortin, J.E. (2013) External conditions inversely change the RNA polymerase II elongation rate and density in yeast. Biochim Biophys Acta, 1829, 1248-1255.

55. Pelechano, V., Jimeno-Gonzalez, S., Rodriguez-Gil, A., Garcia-Martinez, J., Perez-Ortin, J.E. and Chavez, S. (2009) Regulon-specific control of transcription elongation across the yeast genome. PLoS Genet, 5, e1000614.

56. Barnes, C.O., Calero, M., Malik, I., Graham, B.W., Spahr, H., Lin, G., Cohen, A.E., Brown, I.S., Zhang, Q., Pullara, F. et al. (2015) Crystal Structure of a Transcribing RNA Polymerase II Complex Reveals a Complete Transcription Bubble. Mol Cell, 59, 258-269.

57. Millan-Zambrano, G., Rodriguez-Gil, A., Penate, X., de Miguel-Jimenez, L., Morillo-Huesca, M., Krogan, N. and Chavez, S. (2013) The prefoldin complex regulates chromatin dynamics during transcription elongation. PLoS Genet, 9, e1003776.

58. Kim, M., Krogan, N.J., Vasiljeva, L., Rando, O.J., Nedea, E., Greenblatt, J.F. and Buratowski, S. (2004) The yeast Rat1 exonuclease promotes transcription termination by RNA polymerase II. Nature, 432, 517-522.

59. Luo, W., Johnson, A.W. and Bentley, D.L. (2006) The role of Rat1 in coupling mRNA 3’-end processing to transcription termination: implications for a unified allosteric-torpedo model. Genes Dev, 20, 954-965.

60. Jimeno-Gonzalez, S., Schmid, M., Malagon, F., Haaning, L.L. and Jensen, T.H. (2014) Rat1p maintains RNA polymerase II CTD phosphorylation balance. RNA, 20, 551-558.

61. Jiao, X., Xiang, S., Oh, C., Martin, C.E., Tong, L. and Kiledjian, M. (2010) Identification of a quality-control mechanism for mRNA 5’-end capping. Nature, 467, 608-611.

62. Chang, J.H., Jiao, X., Chiba, K., Oh, C., Martin, C.E., Kiledjian, M. and Tong, L. (2012) Dxo1 is a new type of eukaryotic enzyme with both decapping and 5’-3’ exoribonuclease activity. Nat Struct Mol Biol, 19, 1011-1017.

63. Chen, H., Shiroguchi, K., Ge, H. and Xie, X.S. (2015) Genome-wide study of mRNA degradation and transcript elongation in Escherichia coli. Mol Syst Biol, 11, 808.

64. Nehlin, J.O., Carlberg, M. and Ronne, H. (1991) Control of yeast GAL genes by MIG1 repressor: a transcriptional cascade in the glucose response. EMBO J, 10, 3373-3377.

65. Gancedo, J.M. (1998) Yeast carbon catabolite repression. Microbiol Mol Biol Rev, 62, 334-361.

66. De Vit, M.J., Waddle, J.A. and Johnston, M. (1997) Regulated nuclear translocation of the Mig1 glucose repressor. Mol Biol Cell, 8, 1603-1618.

67. Shaw, R.J., Wilson, J.L., Smith, K.T. and Reines, D. (2001) Regulation of an IMP dehydrogenase gene and its overexpression in drug-sensitive transcription elongation mutants of yeast. J Biol Chem, 276, 32905-32916.

68. Hyle, J.W., Shaw, R.J. and Reines, D. (2003) Functional distinctions between IMP dehydrogenase genes in providing mycophenolate resistance and guanine prototrophy to yeast. J Biol Chem, 278, 28470-28478.

69. Jenks, M.H. and Reines, D. (2005) Dissection of the molecular basis of mycophenolate resistance in Saccharomyces cerevisiae. Yeast, 22, 1181-1190.

70. Exinger, F. and Lacroute, F. (1992) 6-Azauracil inhibition of GTP biosynthesis in Saccharomyces cerevisiae. Curr Genet, 22, 9-11.

71. Hartzog, G.A., Wada, T., Handa, H. and Winston, F. (1998) Evidence that Spt4, Spt5, and Spt6 control transcription elongation by RNA polymerase II in Saccharomyces cerevisiae. Genes Dev, 12, 357-369.

72. Gaillard, H., Tous, C., Botet, J., Gonzalez-Aguilera, C., Quintero, M.J., Viladevall, L., Garcia-Rubio, M.L., Rodriguez-Gil, A., Marin, A., Arino, J. et al. (2009) Genome-wide analysis of factors affecting transcription elongation and DNA repair: a new role for PAF and Ccr4-not in transcription-coupled repair. PLoS Genet, 5, e1000364.

73. Gaur, N.A., Hasek, J., Brickner, D.G., Qiu, H., Zhang, F., Wong, C.M., Malcova, I., Vasicova, P., Brickner, J.H. and Hinnebusch, A.G. (2013) Vps factors are required for efficient transcription elongation in budding yeast. Genetics, 193, 829-851.

74. Squazzo, S.L., Costa, P.J., Lindstrom, D.L., Kumer, K.E., Simic, R., Jennings, J.L., Link, A.J., Arndt, K.M. and Hartzog, G.A. (2002) The Paf1 complex physically and functionally associates with transcription elongation factors in vivo. EMBO J, 21, 1764-1774.

75. Denis, C.L., Chiang, Y.C., Cui, Y. and Chen, J. (2001) Genetic evidence supports a role for the yeast CCR4-NOT complex in transcriptional elongation. Genetics, 158, 627-634.

76. Davis, C.A. and Ares, M., Jr. (2006) Accumulation of unstable promoter-associated transcripts upon loss of the nuclear exosome subunit Rrp6p in Saccharomyces cerevisiae. Proc Natl Acad Sci U S A, 103, 3262-3267.

77. Steinmetz, E.J., Warren, C.L., Kuehner, J.N., Panbehi, B., Ansari, A.Z. and Brow, D.A. (2006) Genome-wide distribution of yeast RNA polymerase II and its control by Sen1 helicase. Mol Cell, 24, 735-746.

78. Canadell, D., Garcia-Martinez, J., Alepuz, P., Perez-Ortin, J.E. and Arino, J. (2015) Impact of high pH stress on yeast gene expression: A comprehensive analysis of mRNA turnover during stress responses. Biochim Biophys Acta, 1849, 653-664.

79. Howard, S.C., Chang, Y.W., Budovskaya, Y.V. and Herman, P.K. (2001) The Ras/PKA signaling pathway of Saccharomyces cerevisiae exhibits a functional interaction with the Sin4p complex of the RNA polymerase II holoenzyme. Genetics, 159, 77-89.

80. Howard, S.C., Hester, A. and Herman, P.K. (2003) The Ras/PKA signaling pathway may control RNA polymerase II elongation via the Spt4p/Spt5p complex in Saccharomyces cerevisiae. Genetics, 165, 1059-1070.

81. Darby, M.M., Serebreni, L., Pan, X., Boeke, J.D. and Corden, J.L. (2012) The Saccharomyces cerevisiae Nrd1-Nab3 transcription termination pathway acts in opposition to Ras signaling and mediates response to nutrient depletion. Mol Cell Biol, 32, 1762-1775.

82. Cui, P., Jin, H., Vutukuru, M.R. and Kaplan, C.D. (2016) Relationships Between RNA Polymerase II Activity and Spt Elongation Factors to Spt - Phenotype and Growth in Saccharomyces cerevisiae. G3 (Bethesda), 6, 2489-2504.

83. Rasmussen, E.B. and Lis, J.T. (1993) In vivo transcriptional pausing and cap formation on three Drosophila heat shock genes. Proc Natl Acad Sci U S A, 90, 7923-7927.

84. Glover-Cutter, K., Kim, S., Espinosa, J. and Bentley, D.L. (2008) RNA polymerase II pauses and associates with pre-mRNA processing factors at both ends of genes. Nat Struct Mol Biol, 15, 71-78.

85. Hu, W., Petzold, C., Coller, J. and Baker, K.E. (2010) Nonsense-mediated mRNA decapping occurs on polyribosomes in Saccharomyces cerevisiae. Nat Struct Mol Biol, 17, 244-247.

86. Hu, W., Sweet, T.J., Chamnongpol, S., Baker, K.E. and Coller, J. (2009) Co-translational mRNA decay in Saccharomyces cerevisiae. Nature, 461, 225-229.

87. Pelechano, V., Wei, W. and Steinmetz, L.M. (2015) Widespread Co-translational RNA Decay Reveals Ribosome Dynamics. Cell, 161, 1400-1412.

88. Larson, D.R., Zenklusen, D., Wu, B., Chao, J.A. and Singer, R.H. (2011) Real-time observation of transcription initiation and elongation on an endogenous yeast gene. Science, 332, 475-478.

89. Hocine, S., Raymond, P., Zenklusen, D., Chao, J.A. and Singer, R.H. (2013) Single-molecule analysis of gene expression using two-color RNA labeling in live yeast. Nat Methods, 10, 119-121.

90. Mason, P.B. and Struhl, K. (2003) The FACT complex travels with elongating RNA polymerase II and is important for the fidelity of transcriptional initiation in vivo. Mol Cell Biol, 23, 8323-8333.

